# Aiki-GeNano: Multi-Stage Preference Optimization for Generative Design of Developable Nanobodies

**DOI:** 10.64898/2026.04.28.721526

**Authors:** Radheesh Sharma Meda, Jigar Doshi, Eswar Iyer, Shankar Shastry, Venkatesh Mysore

## Abstract

Therapeutic nanobodies must combine target binding with biophysical and chemical properties that determine manufacturability, stability, and clinical viability, collectively termed developability, yet most computational design pipelines still treat developability as a post-hoc filter rather than an integrated training objective. We present Aiki-GeNano, a three-stage language-model alignment pipeline for epitope-conditioned nanobody generation that integrates multiple developability signals directly into training, using only sequence information and previously published predictors. Across 65 target epitopes and relative to the supervised baseline, the combined pipeline raised predicted mean melting temperature by 6.6 °C, halved isomerization-motif severity, reduced deamidation, N-glycosylation sequons and CDR methionine-oxidation motifs, and preserved predicted humanness and solubility. On a shared 10-target GPCR benchmark, Aiki-GeNano achieved the highest predicted melting temperature and the lowest isomerization severity among five contemporary VHH generators. Starting from ProtGPT2 and a 1.35-million-pair binder dataset generated on an mRNA-display platform, the pipeline applies supervised fine-tuning, Direct Preference Optimization on 522,800 pairs ranked by a composite of selectivity, predicted thermal stability, solubility, and humanness, and Group Reward-Decoupled Policy Optimization against six sequence-based rewards (FR2 hydrophobicity, hydrophobic-patch coverage, chemical-liability motifs, Wilkinson–Harrison expression probability, VHH hallmark residues, scaffold integrity). Generated sequences differ from the nearest training sequence by a mean of 8.1–9.0 amino acids out of 126, and two alternative training trajectories converge to distinct amino-acid-composition strategies with similar liability outcomes but different thermal-stability gains, indicating initialization-dependent convergence of the reward-optimized policy. Predicted humanness was preserved at the level of the camelid VHH scaffold of the training library — a data-side limitation rather than a methodological one, since the framework was effectively constant across all preference pairs. Applicability to the drug discovery and development pipeline, limitations of predicted-property evaluation, and future work are discussed.

## 1 Introduction

Nanobodies are single-domain antibody fragments derived from the heavy-chain-only immunoglobulins of camelids (VHH) and sharks (VNAR). Owing to their small size (∼ 15 kDa), high thermal stability, deep-tissue penetration, and amenability to microbial expression, they offer several advantages over conventional antibodies as a therapeutic modality [1]. A principal concern has been their non-human origin and the associated risk of immunogenicity in patients; however, advances in humanization and engineering have progressively mitigated this barrier, culminating in four approved nanobody-based drugs: caplacizumab for acquired thrombotic thrombocytopenic purpura (2018), envafolimab for advanced solid tumors (2021), ciltacabtagene autoleucel as a CAR-T therapy for multiple myeloma (2022), and ozoralizumab for rheumatoid arthritis (2022) [2–5].

Like full-length antibodies (IgGs), the primary mechanism of action is highly specific binding of a portion of the nanobody surface (paratope) to a region of the therapeutic target (epitope) through a broad protein-protein interface. The three complementarity-determining regions (CDRs 1, 2, and 3) of the single VHH domain dominate this interaction, interweaving with four framework regions (FRs 1–4). This binding either directly elicits the functional response or serves as a targeted delivery vehicle for additional therapeutic payloads. In both cases, the binding affinity and selectivity of the interface are the defining identities of a hit- or lead-stage binder. By the time a candidate progresses to drug development, several additional considerations surface that collectively determine whether a reasonable path exists to a viable human therapeutic. The contributing factors specific to protein biologics are collectively termed *developability* and span solubility, aggregation propensity, expression yield, thermal stability, proteolytic stability, clearance, half-life, and immunogenicity. Chemical instability, primarily asparagine deamidation and aspartate isomerization, has been recognized since the late 1990s as accounting for 40–50% of formulation failures in protein therapeutics [6], with aggregation causing an additional 30–35% of manufacturing losses [7]. Systematic analysis of 137 clinical-stage antibodies confirmed chemical degradation as the predominant liability [8].

Developability evaluation remains predominantly a post-hoc filtering step rather than an integrated design objective [9]. Despite representing ∼$1–2 billion in sunk investment per failed program [10], developability-driven attrition has persisted for three broad reasons. First, several relevant properties (e.g., immunogenicity) are inherently difficult to predict confidently, and the predictions are accordingly less actionable. Second, improving developability often worsens binding: for example, affinity-driven mutagenesis may introduce sequence motifs such as asparagine-glycine (deamidation half-life 1–3 days at pH 7.4, 37 ◦C [11]) or aspartate-glycine / aspartate-serine (isomerization half-life 30–90 days [12]), both predicted chemical liabilities. Third, the consequences of non-ideal developability properties tend to manifest late in the drug’s development journey when fewer alternative candidates remain. Unlike small molecules, for which ADMET frameworks provide systematic early-stage screens [13], therapeutic biologics lack equivalent validated computational pipelines that integrate reliable multi-property optimization into the generative process itself. Nanobodies face further scaffold-specific considerations [1, 14], notably a solvent-exposed framework region 2 (FR2) in the absence of a light-chain (V_L_) partner, making standard full-length-antibody developability tools only partially applicable.

Computational approaches to nanobody design have advanced considerably but remain fragmented in their treatment of developability. Raybould *et al*. codified five computational guidelines for therapeutic antibody profiling (CDR length, surface hydrophobicity, CDR positive and negative charge, and asymmetry in heavy/light-chain surface charges) that serve as downstream filters [15]. Structure-based generative methods including RFdiffusion [16] and RFantibody [17] generate structurally plausible CDR conformations but require solved or predicted co-complex structures as input and do not optimize for chemical developability during generation. Sequence-based protein language models such as ProtGPT2 [18], IgLM [19], and PepMLM [20] enable scalable epitope-conditioned generation but do not support developability as a training objective, leaving the generative model blind to biophysical and biochemical processes that cause late-stage failures.

Several techniques have recently been proposed for incorporating developability into the generative process directly, each with limitations for our use case. Guided discrete diffusion for conventional IgG antibodies [21, 22] steers generation toward improved hydrophobicity and reduced self-association, but relies on regression oracles trained on quantitative assay measurements from clinical-stage antibodies and is not directly transferable to nanobodies. Our prior work on preference optimization of protein language models [23] demonstrated DPO-driven alignment for peptide generation at small scale (∼ 9,000 pairs) with a single physicochemical objective (isoelectric point). Other recent DPO applications to protein engineering include ProteinDPO [24], which demonstrated how structural stability can be imparted onto a generative model using DPO over mutational data; ProteinMPNN preference optimization for immunogenicity reduction [25]; and scalable DPO variants for protein language models evaluated on thermostability and expression mutational datasets [26]. Related reinforcement-learning-style peptide design efforts have explored preference optimization and structural inference [27, 28]. Nanobody-specific efforts have begun to close this gap on the scoring and generation sides: nanoBERT [29] provides a VHH-specific masked language model; NanoAbLLaMA [30] and IgGM [31] support scaffolded generation; a nativeness-constrained diffusion framework [32] enforces an evolutionary prior at inference time; and the Therapeutic Nanobody Profiler [33] and AbNatiV [34] provide nanobody-calibrated developability and nativeness scoring. Epitope-conditioned generation with reward-level optimization across the multi-property developability spectrum considered here has not, to our knowledge, been demonstrated before. Further, no existing generative protein-design framework that applies preference-based alignment addresses the reward-signal collapse that arises when combining objectives with heterogeneous variance, a problem formalized for multi-reward reinforcement learning by Liu *et al*. [35].

Here we present a three-stage language-model training framework, termed Aiki-GeNano, that directly encodes multi-property developability into epitope-conditioned nanobody generation. Starting from the generative protein language model ProtGPT2 [18], we perform supervised fine-tuning (SFT) on 1.35 million nanobody– epitope pairs across 65 diverse target epitopes to establish binding-compatible sequence priors, followed by Direct Preference Optimization (DPO) on 522,800 preference pairs ranked by a composite developability score integrating thermal stability, solubility, and humanness. As a third stage, we apply Group Reward-Decoupled Policy Optimization (GDPO) [35], which resolves multi-reward signal collapse by normalizing each of six mechanistic reward signals independently before the policy update, with a KL divergence term that anchors the policy to the initialization distribution (see Eq. 6). The rewards were designed from established biophysical literature: (R_1_) FR2-specific hydrophobicity penalizing the solvent-exposed framework region unique to nanobodies [1, 36]; (R_2_) a hydrophobic-patch reward identifying consecutive aggregation-prone residue runs inspired by AGGRESCAN [37]; (R_3_) chemical liability scoring deamidation-prone dipeptides weighted by half-life severity [11, 38], isomerization-risk motifs [12, 39], fragmentation sites, N-glycosylation sequons, and CDR methionine oxidation, carrying the highest reward weight; (R_4_) expression probability predicting soluble *E. coli* expression from sequence composition [40]; and (R_5_) VHH hallmark conservation enforcing hydrophilic substitutions at the four FR2 tetrad positions required in the absence of a V_L_ partner [1]. Collectively, these objectives build developability assessment into training rather than into a post-hoc filter.

## 2 Results

Preliminary screening data involving a 126-residue VHH library (see *Materials and Methods*) was treated as ground-truth binding data for the computational experiments reported here; the focus is on conferring developability properties to target-specific distributions of binder sequences (rather than improving binding affinity). All developability properties were computed using pre-existing computational tools or published heuristics that require only protein sequence information (see *Supplementary Information*). While the properties were chosen to be as broad as pragmatically possible, they are meant to serve as a representative set rather than a complete recipe. An overview of the training pipeline, the four DPO preference axes (selectivity, *T*_*m*_, solubility, humanness), the six GDPO reward functions, and the VHH scaffold regions each reward acts on is given in Figure 1.

**Fig. 1.**
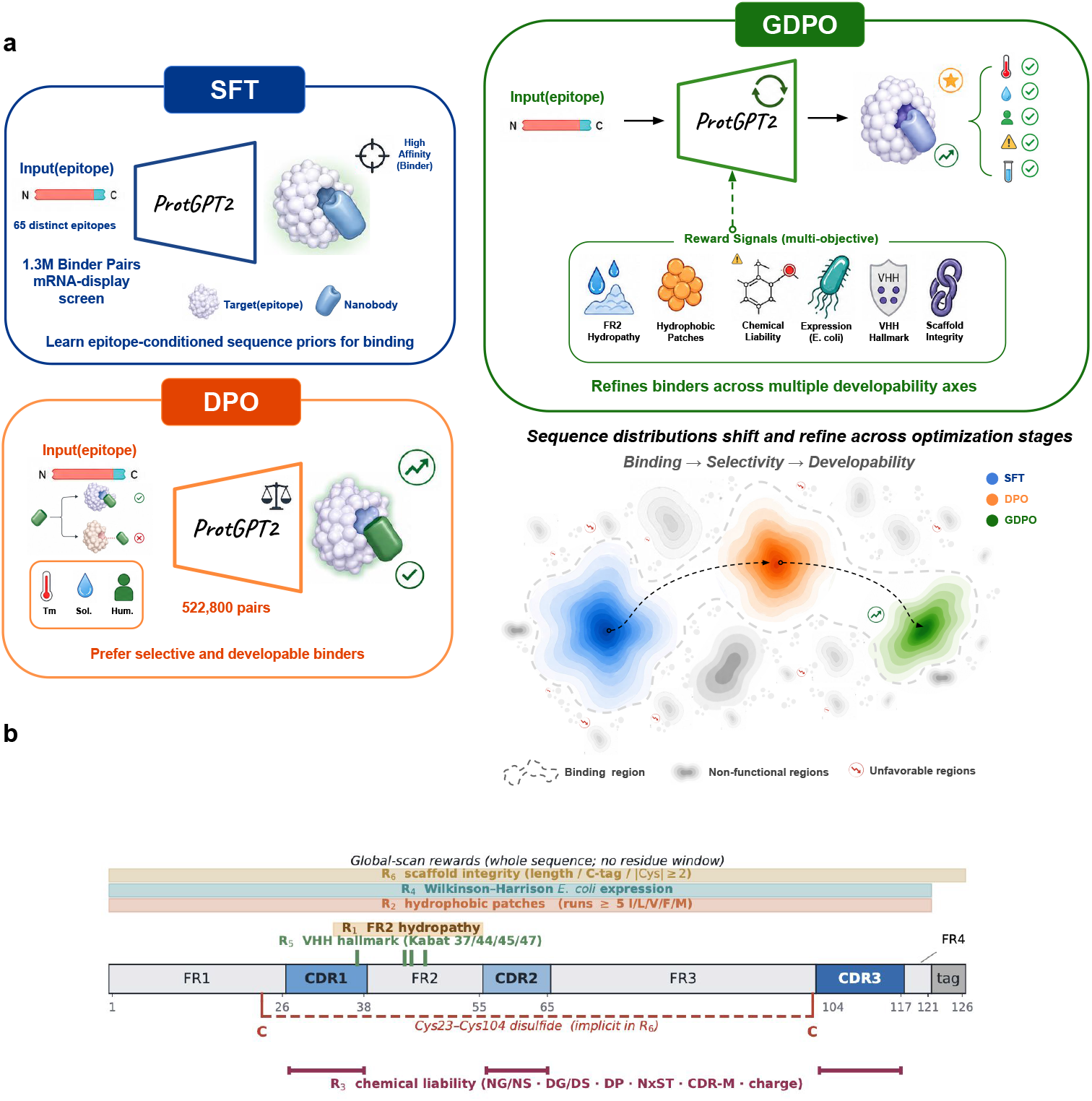
**a**, Three-stage alignment of ProtGPT2 for epitope-conditioned nanobody generation. Stage 1 (SFT): supervised fine-tuning on 1.35 M nanobody–epitope pairs from an mRNA-display screen across 65 targets establishes a binding-conditioned sequence prior. Stage 2 (DPO): Direct Preference Optimization on 522,800 pairs ranked by selectivity, predicted thermal stability (Tm), solubility, and humanness aligns the model with global developability preferences. Stage 3 (GDPO): Group Reward-Decoupled Policy Optimization refines the model against six mechanistic reward signals with per-reward normalization, while retaining DPO-stage properties. Aiki-GeNano represents the GDPO path initialized from the DPO checkpoint (SFT → DPO → GDPO); a second pathway directly from SFT (SFT → GDPO) is also evaluated. The density landscape (lower right) illustrates how the two training trajectories explore distinct regions of developability space: GDPO(DPO) (green) and GDPO(SFT) (red) converge on different compositional strategies despite optimizing identical reward functions, with arrows indicating the direction of progressive optimization from the SFT baseline (blue). **b**, Sequence-level map of where each reward function acts on the 126-residue VHH scaffold. *R*_1_ penalizes mean hydrophobicity across FR2 (positions 36–53); *R*_5_ scores the Kabat VHH hallmark tetrad (37, 44, 45, 47); *R*_2_ and *R*_4_ act over the full 121-residue core sequence; *R*_3_ scans CDR regions for deamidation (NG/NS), isomerization (DG/DS), fragmentation (DP), N-glycosylation (NxST), CDR methionine oxidation, and charge clusters; *R*_6_ enforces length, C-terminal tag, and the canonical Cys23–Cys104 disulfide.

We trained conditional nanobody binder models through three sequential stages:

- Supervised fine-tuning (SFT) on native nanobody–target pairs from the raw screening data.
- Direct preference optimization (DPO) along four developability axes (selectivity, melting temperature, solubility, and humanness), using pairs of binders for a target in which one exhibited distinctly poorer properties. Binders to a target were paired with binders to dissimilar targets as their less-preferred counterparts, to reinforce target specificity.
- Group reward-decoupled policy optimization (GDPO) to maximize a composite reward function comprising six developability objectives: FR2 aggregation propensity minimization, hydrophobic patch curtailment, chemical liability reduction (deamidation, isomerization, N-glycosylation, oxidation), expression-yield maximization, VHH hallmark motif preservation, and scaffold integrity. A KL term penalized deviation of the generated distribution from the initialization distribution.

Two GDPO variants were explored: one initialized from the DPO checkpoint (SFT → DPO → GDPO, termed Aiki-GeNano) and one initialized directly from SFT (SFT → GDPO), allowing us to probe how the initialization influences the final property distributions when both paths optimize identical reward functions. Throughout, “Aiki-GeNano family” refers collectively to the four training-stage models (SFT, DPO, GDPO(SFT), and GDPO(DPO)) and “Aiki-GeNano” specifically to the GDPO(DPO) checkpoint. All models were evaluated by generating 100 sequences per target (sampling temperature *T* = 0.7, nucleus sampling top-*p* = 0.9) conditioned on a single target epitope (peptide, IDR, or folded domain) and assessed developability across five biophysical properties: thermal stability, chemical stability, expression, structural stability, and framework conservation.

### 2.1 DPO adds selectivity and early-stage developability

Supervised fine-tuning teaches the model to generate binders for a given target epitope but provides no guidance on which binder sequences to avoid. The DPO stage addresses two drug development concerns immediately after binding: *selectivity* (the same paratope should not also bind unrelated targets) and *broad early-stage developability* (predicted thermal stability, solubility and humanness, the three axes most often used for first-pass triage). Selectivity is encouraged by treating binders to sequence-distant targets as dispreferred examples (a quantitative selectivity readout on the trained model is outside the scope of this paper and is identified as future work), and developability by ranking within-target pairs on a composite score of predicted thermal stability, solubility, and humanness (Eq. 1). DPO operates on 522,800 such pairs and shifts the prior accordingly: at the global property level, DPO achieves a mean thermal stability gain of +3.0 ^◦^C over the SFT baseline (SFT: 71.7 ^◦^C; DPO: 74.7 ^◦^C, *T* = 0.7, *n* = 3 seeds), accompanied by a reduction in instability index (36.1 → 34.3) and an increase in predicted solubility of +0.016 NetSolP units, all while fully preserving sequence humanness (Δ = +0.007).

At the per-motif liability level, however, the behavior is more nuanced. Reductions are limited across most motifs (NS decreases from 0.468 to 0.266; NT and NN remain largely unchanged), while NG deamidation severity *increases* from 0.651 to 1.034. This counterintuitive shift arises because the preference pairs cover only a subset of the developability criteria, forcing the model to infer the remainder indirectly. For instance, the model can incorrectly learn to favor sequences that incidentally carry a higher NG motif frequency at certain CDR positions. DPO is too coarse to penalize those localized liabilities and instead reinforces them through global preference alignment. Correcting these localized chemical vulnerabilities without sacrificing the binding, selectivity, and broad developability gains already established by DPO requires a fundamentally different optimization strategy: one that can resolve individual liability motifs as independent objectives. This is precisely the role of the GDPO stage.

### 2.2 GDPO adds the per-property developability considerations that DPO’s composite signal cannot capture

To evaluate how GDPO builds on the foundation established by DPO, we assessed aggregate developability performance across all generated sequences from each training stage (Figure 2). The radar plot (Fig. 2a) revealed distinct optimization trajectories for the two GDPO variants: the SFT → DPO → GDPO path achieved the strongest gains in thermal stability and chemical stability, while the direct SFT → GDPO path showed more balanced but smaller improvements than the Aiki-GeNano path. Both GDPO variants outperformed their respective baselines on the composite developability profile. Thermal stability, quantified as predicted melting temperature (T_*m*_) via TEMPRO, increased progressively across training stages (Fig. 2b). The SFT → DPO → GDPO path achieved a mean T_*m*_ of 78.3 ◦C, representing a +6.6 ◦C improvement over the SFT baseline (71.7 ◦C) and a +3.6 ◦C gain beyond DPO alone (74.7 ◦C). The SFT → GDPO path reached 75.9 ◦C, exceeding DPO by +1.2 ◦C but remaining 2.4 ◦C below the GDPO(DPO) benchmark. Both routes improve predicted thermal stability, with the SFT → DPO → GDPO trajectory yielding the larger improvement. Humanness scores, assessed via the Sapiens antibody language model [41], remained stable across all stages (0.648–0.657; Fig. 2c), indicating that reward-driven optimization preserved the antibody germline characteristics relevant to therapeutic development. Predicted solubility (NetSolP [42]) remained consistent across all stages (0.611–0.635; Fig. 2e), confirming that the developability gains introduced by DPO and GDPO did not come at the expense of solution behavior.

**Fig. 2.**
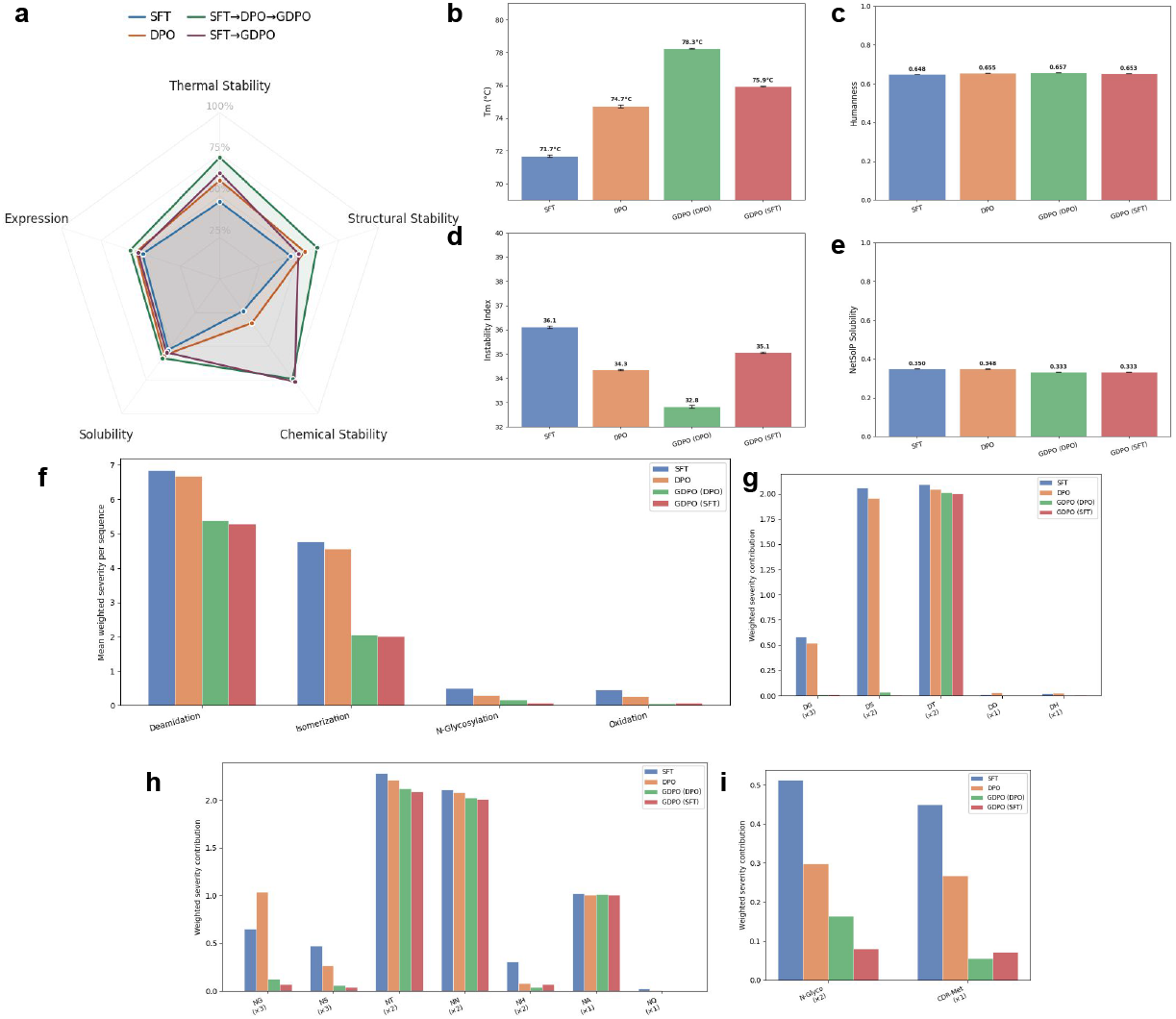
**a**, Radar plot of normalized developability scores across training stages: SFT (base), DPO (preference-aligned), and two GDPO variants: SFT → DPO → GDPO (Aiki-GeNano) and SFT → GDPO. **b**, Mean predicted T_*m*_ increases by +6.6 ^*◦*^C from SFT to DPO GDPO, the best training path by this metric. **c**, All training stages maintain humanness scores around 0.65. **d**, GDPO achieves the lowest mean instability index. **e**, Mean NetSolP solubility remains comparable across all stages. **f**, Liability severity breakdown by type shows that GDPO stages substantially reduce deamidation and isomerization motifs relative to SFT and DPO baselines. **g–i**, Per-position liability contributions across CDR1, CDR2, and CDR3 regions reveal targeted reduction at high-severity sites. All metrics were computed on binders generated at *T* = 0.7 across 65 targets.

Structural stability, quantified via the instability index [43] (lower values indicate greater stability), improved markedly in the SFT → DPO → GDPO path (Fig. 2d), achieving a mean of 32.8 compared to 36.1 for SFT, a reduction of 3.3 units. This improvement occurred despite the instability index not being explicitly included as a reward signal, implying that the motif-level and compositional changes drive a secondary improvement in instability index. GDPO(SFT) maintained an instability index (35.1) comparable to the SFT baseline, confirming that the SFT → DPO → GDPO trajectory more effectively captures this property. Chemical-liability profiles confirmed that reward-based optimization reduced the two dominant liability categories (Fig. 2f–i). Both GDPO variants substantially reduced deamidation and isomerization severity. Deamidation liability decreased from 6.8 (SFT) to 5.4 (GDPO(DPO)) and 5.3 (GDPO(SFT)), while isomerization dropped from 4.7 to 2.0 and 2.0, respectively. The per-position analysis (Fig. 2g–i) revealed that this reduction was spatially concentrated in the CDR regions, particularly at high-severity hotspots (CDR1 NG/NS motifs; CDR2 DG motifs). GDPO models eliminated these hotspot motifs while also reducing N-glycosylation (0.5 → 0.2) and CDR methionine oxidation (0.5 → ∼ 0.0), covering both in addition to the two dominant failure modes.

### 2.3 CDR-localized sequence modifications drive property improvements

To localise these changes within the nanobody sequence and check that the scaffold is preserved, we performed positional frequency analysis and entropy profiling across the 126-residue sequence (Figure 3). Shannon entropy profiles (Fig. 3a) confirmed that sequence variability concentrates in the three CDR regions, with CDR1 (positions 27–38) and CDR3 (positions 105–117) exhibiting the highest entropy (*H >* 2.0 bits at several positions), while framework regions remain nearly invariant (*H <* 0.5 bits). All models preserve this CDR-focused variability pattern, confirming that none of the training stages corrupts the conserved nanobody scaffold (at the recommended sampling temperature, *T* = 0.7; framework variability at higher temperatures is reported in the Supplementary Information).

**Fig. 3.**
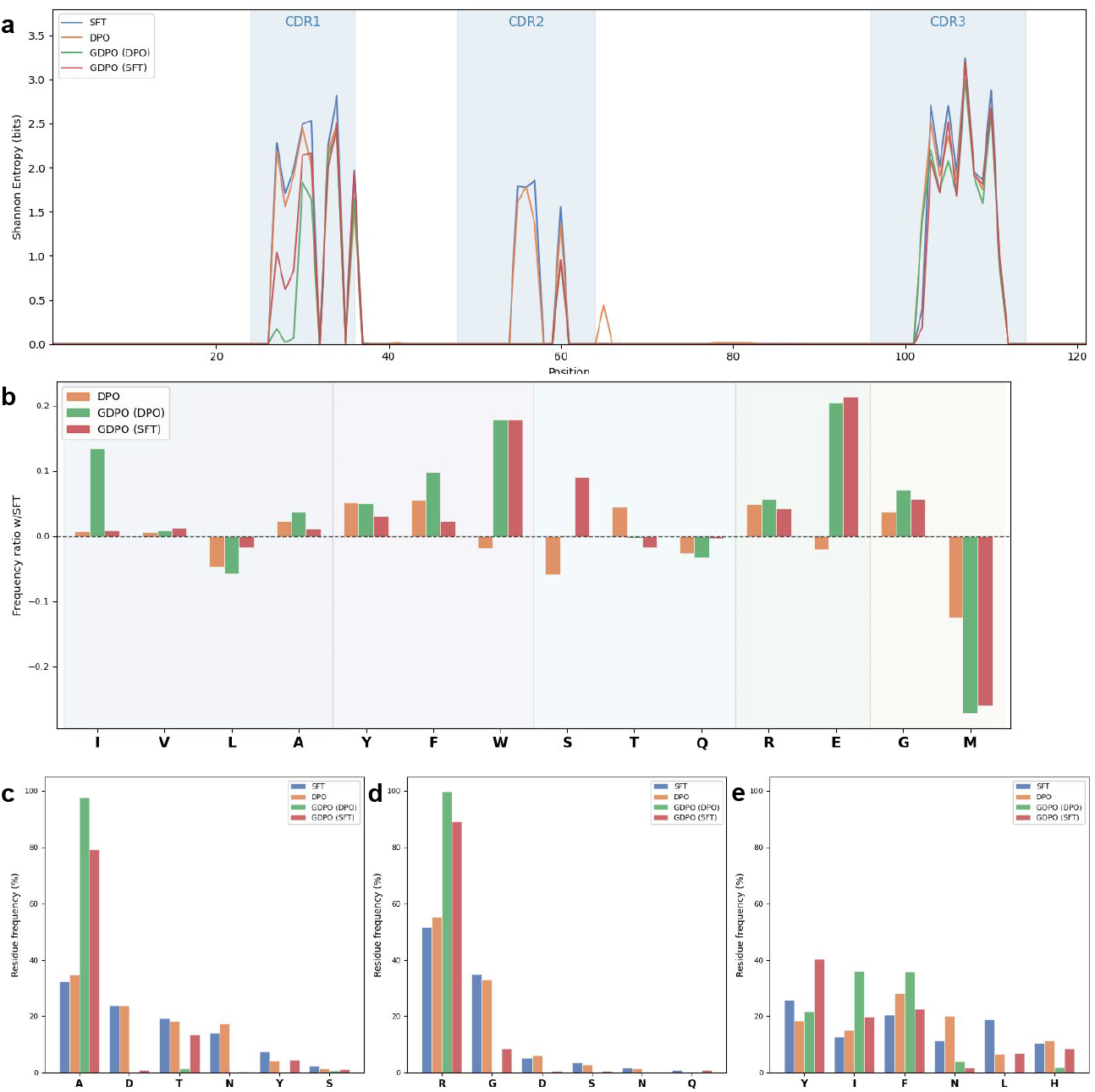
**a**, Positional Shannon entropy across the 126-residue nanobody sequence reveals variability concentrated in CDR regions (CDR1: 27–38; CDR2: 56–65; CDR3: 105–117), while framework regions remain highly conserved across all training stages. **b**, Global amino acid frequency ratio relative to SFT, positive values indicate enrichment, and negative values indicate depletion. Both GDPO variants deplete liability-prone aspartate, asparagine, and methionine while enriching tryptophan; GDPO (DPO) additionally shows strong isoleucine enrichment absent from GDPO(SFT), reflecting path-dependent profile. **c–e**, Residue-frequency distributions at three CDR1 positions: (c) position 27 shows D → A shift in GDPO models (∼ 30% to ∼97%); (d) position 28 shows R enrichment (∼ 55% to ∼98%); (e) position 30 shows N → I/F substitution (∼ 25% to *<*5% N, with I/F increasing). These targeted substitutions eliminate deamidation and isomerization hotspots.

Global amino acid composition analysis (Fig. 3b) reveals three classes of change relative to SFT. First, both GDPO variants converge on shared reward-driven substitutions regardless of initialization: tryptophan is enriched (frequency ratio +0.18 relative to SFT, identical for both paths), while liability-prone residues are coordinately depleted: aspartate (− 0.37/ −0.43; GDPO(DPO)/GDPO(SFT) throughout), asparagine (− 0.17/− 0.19), and methionine (− 0.27/−0.26), consistent with simultaneous optimization of thermal stability and chemical-liability removal. DPO alone does not drive these depletion patterns, confirming GDPO as the stage responsible for liability correction. Second, GDPO(DPO) shows a path-specific isoleucine enrichment (ratio: +0.14) entirely absent from the SFT-initialized variant (+0.01), indicating that DPO pre-training shifts the prior into a region where isoleucine enrichment is favored by the GDPO reward. Third, some DPO-induced compositional shifts are transient and do not survive GDPO optimization: DPO depletes serine (− 0.06 vs. SFT) and enriches threonine (+0.04), but both shifts are suppressed or reversed in the subsequent GDPO stage regardless of initialization path, indicating that GDPO overrides prior compositional changes that are misaligned with the downstream reward.

To identify the precise positions driving these changes, we examined per-position residue frequencies at key CDR1, CDR2, and CDR3 sites (Fig. S3). In CDR1, three positions reveal systematic liability-reducing substitutions (Fig. 3c–e). At position 27, both GDPO variants nearly eliminated aspartate (D: 24% in SFT → ∼1% in GDPO), replacing it predominantly with alanine (A: 32%→98% GDPO(DPO), 79% GDPO(SFT)), removing a high-risk DG/DS isomerization motif while preserving structural compatibility. At position 28, both GDPO models converged on arginine (R: 52%→100% GDPO(DPO), 89% GDPO(SFT)), displacing glycine (G: 35%→*<*1%/*<*9%), which may improve loop stability and binding electrostatics. At position 30, GDPO markedly reduced asparagine (N: 11% in SFT→4% GDPO(DPO), ∼2% GDPO(SFT)); notably, DPO had itself increased N at this position (11% → 20%), inadvertently introducing a deamidation liability that GDPO subsequently corrected. The replacement preferences diverge by path: GDPO(DPO) strongly enriches isoleucine and phenylalanine (I+F: 34% SFT → 72%), while GDPO(SFT) retains more tyrosine (Y: 26% → 40%) alongside moderate I/F enrichment (43%). This position-level I/F enrichment is the positional origin of the path-specific global isoleucine enrichment described before.

Extending the analysis to CDR2 reveals further convergent and divergent patterns. Framework-proximal CDR1 positions 24–26 (A-S-G) are invariant at 100% across all models, confirming scaffold preservation. In CDR2, positions 55–57 converge to a near-consensus S-W-G motif in both GDPO variants (position 55: 41% S in SFT → 100%; position 56: 55% W → 100%; position 57: 63% G → 100%), suggesting these positions form a structural hinge fixed by GDPO regardless of initialization. Position 60 shows path-divergent switching (SFT: R:44%/I:42%→I:64%/R:36% under GDPO(DPO) versus R:62%/I:38% under GDPO(SFT)), with the GDPO(DPO) path favoring hydrophobic I and the GDPO(SFT) path retaining charged R, consistent with the divergent global hydrophobicity trajectories.

In CDR3, the framework-proximal anchor position 102 reveals that GDPO(SFT) stringently conserves leucine (L: 95% SFT → 98%), while GDPO(DPO) tolerates greater variability (L: 77%, with V/A substitutions). At the hypervariable tip, DPO introduced aspartate at position 110 (D: *<*5% in SFT → 16% in DPO), creating a new deamidation-prone residue; GDPO(DPO) only partially reduces this DPO-inherited liability (D: 13%), whereas GDPO(SFT) (initialized from the original SFT distribution) maintains near-zero aspartate and instead enriches glycine (G: 18% SFT → 29%). This position-level observation directly corroborates the incomplete liability correction in the GDPO(DPO) path seen at the global level.

These positional and compositional shifts produce measurably different global sequence profiles. GDPO(DPO), driven by its preferential enrichment of hydrophobic residues, reaches a mean GRAVY score of −0.29, a substantial shift from the SFT baseline of −0.36. GDPO(SFT) achieves comparable liability reduction through a more polar compositional strategy, resulting in a moderate shift to −0.31. Despite this divergence, both paths preserve humanness and solubility (Fig. 2c,e) while improving predicted thermal stability (Fig. 2b), confirming that the reward landscape admits multiple viable trajectories toward the developability objective.

### 2.4 Target-level generalization and path-dependent optimization

The aggregate improvements and sequence-level mechanisms described above could in principle be driven by strong gains on a subset of targets; we therefore examined whether these results generalize across all 65 epitopes. Per-target analysis of thermal stability versus chemical liability (Fig. 4a) confirmed that both GDPO variants achieve consistent improvements across all 65 targets, with no target showing regression below the SFT baseline. The scatter plot revealed clear stratification: GDPO(DPO) occupies the highest T_*m*_ region (upper-right), GDPO(SFT) forms an intermediate cluster, and SFT/DPO baselines occupy the lower-left. For any given liability severity level, GDPO(DPO) consistently achieves higher T_*m*_ than GDPO(SFT), indicating that the GDPO(DPO) path more effectively converts liability reduction into thermal-stability gains (also see Supplementary Fig. S4).

**Fig. 4.**
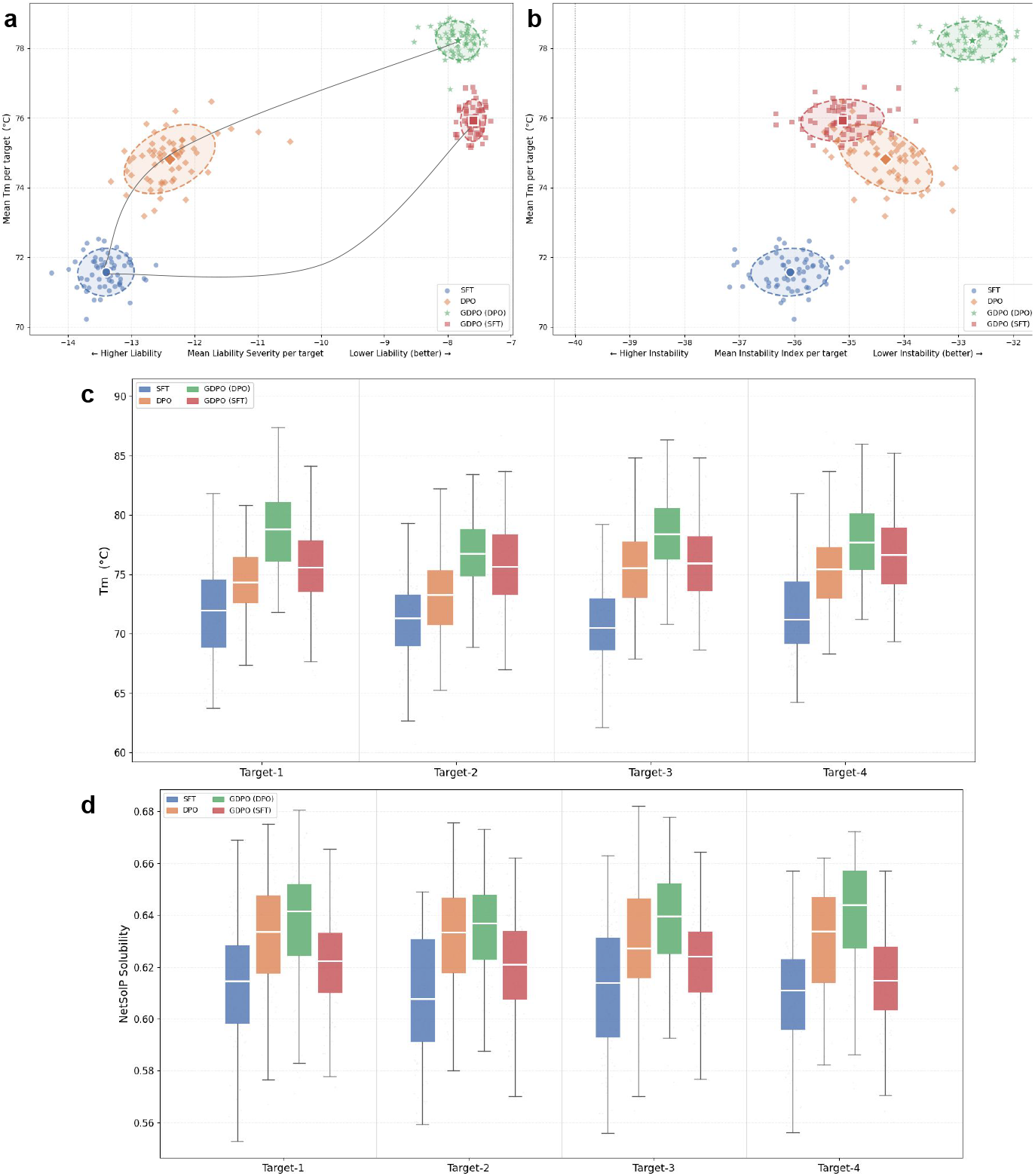
**a**, Per-target mean T_*m*_ vs. mean liability severity across all 65 targets. Both GDPO variants occupy the upper-right quadrant (high T_*m*_, low liability), with GDPO(DPO) achieving consistently higher T_*m*_ than GDPO(SFT) at similar liability levels. **b**, Per-target mean T_*m*_ vs. mean instability index. GDPO(DPO) achieves the strongest separation toward high T_*m*_ and low instability, while GDPO(SFT) shows intermediate improvement. **c**, Predicted T_*m*_ distributions at four representative targets. **d**, NetSolP solubility distributions for the same four targets.

The improvement in structural stability, first observed in aggregate (Fig. 2d), also generalized across targets (Fig. 4b). The T_*m*_ vs. instability-index landscape showed GDPO(DPO) separating strongly toward the upper-right (high T_*m*_, low instability), with minimal overlap with the SFT/DPO baseline clusters. GDPO(SFT) exhibited intermediate performance, achieving higher T_*m*_ than SFT but comparable instability-index values. This path-dependent difference, strong instability reduction under GDPO(DPO) versus modest reduction under GDPO(SFT), suggests that preference alignment installs structural features that compound with subsequent reward optimization. GDPO(SFT), by contrast, reduces liability and lifts T_*m*_ but does not lower the instability index.

To test generalization of the Aiki-GeNano path at the individual target level, four representative targets were selected (Fig. 4c–d): the target where GDPO(DPO) shows the largest dominance margin over all other models; the hardest case where GDPO(DPO) achieves its lowest absolute T_*m*_ across all targets; the target exhibiting the largest T_*m*_ lift from the SFT baseline; and the target where the two GDPO initialization paths produce the most similar distributions, the case most favorable to GDPO(SFT). Across all four representative targets, GDPO(DPO) produced the highest-T_*m*_ sequences with clear separation from DPO and SFT baselines, and the model ranking is preserved. On the most adversarial target, the GDPO(SFT) margin narrows but the ranking does not invert.

Solubility analysis across the same targets (Fig. 4d) confirmed that the same ordering extends beyond thermal stability, with all models preserving comparable developability profiles and GDPO(DPO) maintaining the highest predicted solubility throughout.

### 2.5 Landscape positioning against contemporary nanobody generators

To place Aiki-GeNano in the context of the current generative-nanobody landscape, we re-generated sequences from seven publicly available generators on a shared set of 10 GPCR target epitopes and scored the outputs through the same developability pipeline used for Aiki-GeNano: five VHH-class tools (nanoBERT, IgLM, NanoAbLLaMA, ProteinDPO, IgGM) form our head-to-head set, and two further tools (ProtGPT2, a general-protein LM trained on UniRef50; and PepMLM, designed for 10-AA peptide binders) sit outside the nanobody length range under their designed use cases and are reported only as out-of-class controls. Aiki-GeNano sequences are generated on the fixed 126-residue VHH backbone based on Conrath *et al*. [44]. In contrast, sequences generated by the surveyed tools are on variable nanobody backbones and lengths; hence the canonical 110–130 AA VHH length window is used to filter out sequences that are not nanobodies.

The resulting landscape (Fig. 5; full per-tool *µ*±*σ* table: Supplementary Table S2) shows a pattern of *complementary strengths*. Aiki-GeNano wins decisively on the three axes its DPO and GDPO objectives targeted: the highest mean predicted *T*_*m*_ at 78.3 ◦C, +5 ^◦^C above the best competitor NanoAbLLaMA (73.2 ◦C; pooled-*σ* effect size ≈+1*σ*), the highest NetSolP solubility at 0.635 (+1.5*σ* over NanoAbLLaMA 0.598), and one of the lowest isomerization severities at 2.06 (+1.5*σ* over nanoBERT 4.19; indistinguishable within *σ* from the GDPO(SFT) ablation at 2.02 ± 0.24). *T*_*m*_ and NetSolP solubility were DPO objectives amplified by GDPO; isomerization severity is the direct consequence of GDPO reward *r*_3_. The deamidation severity is lower than for all external tools except IgGM (5.01 vs. 5.39, 0.3*σ*). Conversely, NanoAbLLaMA’s Sapiens humanness (0.750) is ∼9*σ* above Aiki-GeNano’s (0.657), while sequences generated by ProteinDPO show the best instability index (28.9 vs. 32.8 for Aiki-GeNano); the four VHH-class competitors and Aiki-GeNano cluster within 31.6–34.1 on this axis, with ProteinDPO the sole outlier below. One confounding factor is the VHH scaffold being fixed in Aiki-GeNano, restricting humanness gains to CDR substitutions; no alternative frameworks were present in the SFT, DPO or GDPO training data. Additionally, the canonical-VHH content of generated sequences from other tools could not be controlled beyond the length range. A stricter VHH-shape filter (canonical N-terminus and FR4 motif) classifies 100% of nanoBERT and IgGM outputs, 85% of NanoAbLLaMA, 91% of IgLM (the failures concentrated at the N-terminal Q residue), and only 0.6% of ProteinDPO as canonical VHHs; this last observation in particular reframes ProteinDPO’s instability/solubility wins as “novel-protein” wins rather than “nanobody” wins, and each of these confounders is addressed in the *Landscape benchmark scope* subsection of the Limitations.

**Fig. 5.**
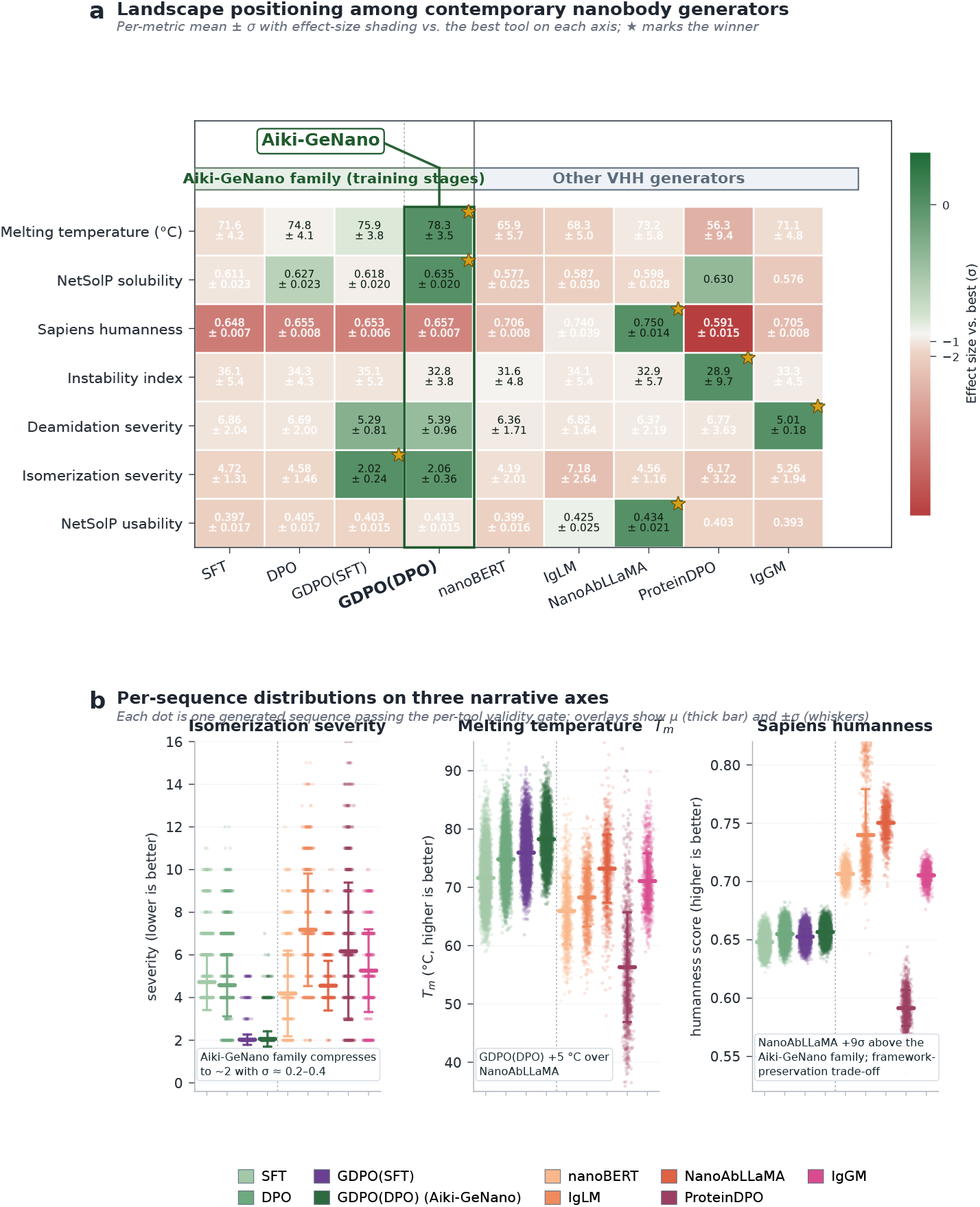
Landscape positioning of Aiki-GeNano against contemporary nanobody generators on a shared 10-target GPCR epitope cohort. Among the four Aiki-GeNano-family columns, only *GDPO(DPO)* (rightmost, starred) is the model referred to as *Aiki-GeNano*; the other three (SFT, DPO, GDPO(SFT)) are training-stage ablations included to expose the contribution of each stage. **a**, Per-metric mean ±*σ* for each tool on eight developability axes, with cell shading indicating effect size (pooled-*σ* units) relative to the best tool on that axis. Stars mark the per-axis winner. Aiki-GeNano-family sequences are generated to a fixed 126-residue VHH backbone (NBv1 scaffold, based on Conrath *et al*. 2001 [44]); competitor sequences are filtered to the canonical VHH length range (110–130 AA, |Cys|≥2). **b**, Per-sequence distributions on three narrative metrics. *Left:* isomerization severity — the single largest win for Aiki-GeNano, whose *r*_3_ reward compresses D-X motif load to *σ* ≈ 0.2–0.4 versus 1.2–3.2 for competitors. *Center:* predicted *T*_*m*_ — GDPO(DPO) sits ∼ 5 ^*◦*^C above the best competitor (NanoAbLLaMA). *Right:* Sapiens humanness — the decisive competitor win, with NanoAbLLaMA ∼ 9*σ* above the Aiki-GeNano family; this trade-off is discussed in the Limitations. Each dot is one generated sequence passing the per-tool validity gate (Aiki: N ≈ 6,400 per model; competitors: N ≈ 1,000). Overlays mark *µ* (bar) and ±*σ* (whiskers).

Per-target rankings across the 10 GPCR epitopes are tool-driven rather than target-driven. Restricting attention to the five non-Aiki VHH-class generators, NanoAbLLaMA leads *T*_*m*_ on 9 of 10 targets (IgGM leads the remaining one) and Sapiens humanness on 10 of 10; ProteinDPO leads the instability index on 10 of 10. Adding Aiki-GeNano to the comparison, GDPO(DPO) leads *T*_*m*_ on 10 of 10 targets. In Pareto terms (humanness vs. *T*_*m*_ and isomerization severity vs. *T*_*m*_), no tool occupies both the high-humanness and the high-*T*_*m*_/low-liability corners simultaneously; NanoAbLLaMA holds the former, Aiki-GeNano (GDPO(DPO)) holds the latter (per-tool *µ*±*σ* in Supplementary Table S2). All five VHH-class comparators emit sequences whose lengths concentrate within the canonical 110–130 AA nanobody window (medians 121–127 AA). Two further tools (ProtGPT2, PepMLM) sit outside the VHH length range under their designed use cases and are omitted from the head-to-head comparison; rationale in Methods §*Landscape benchmark against contemporary generators*.

## 3 Discussion

Mechanistic reward engineering, targeting documented causes of therapeutic failure rather than downstream metric scores, recovers biophysical properties that the GDPO stage never directly optimized. The rewards penalize specific chemical-liability motifs, enforce scaffold constraints, and promote expression-compatible composition, yet predicted T_*m*_ improved by +6.6 ◦C and the instability index decreased by 3.3 units through the SFT → DPO → GDPO path, both emerging as indirect consequences of the underlying sequence changes. The mechanism is interpretable: removing asparagine and aspartate at liability positions eliminates destabilizing polar contributions, and the compensatory enrichment of alanine, isoleucine, and phenylalanine shifts composition toward residues associated with tighter core packing and reduced conformational entropy. Because physicochemical properties are interdependent, decomposing fitness into mechanistically grounded proxy rewards is effective when the decomposition reflects the underlying biophysics, as our results show for nanobody developability. The DPO stage, which explicitly optimizes predicted T_*m*_, solubility, and humanness, likely primes the sequence distribution in ways that amplify these emergent gains under GDPO, consistent with the weaker emergent improvement observed in GDPO(SFT).

Two training trajectories optimizing identical reward functions converge on distinct compositional strategies, with implications beyond nanobody design. GDPO(DPO) enriches hydrophobic residues at key CDR positions (mean GRAVY −0.29) and achieves the largest thermal-stability gains, while GDPO(SFT) retains more polar and aromatic residues (mean GRAVY −0.31) with comparable liability reduction but intermediate T_*m*_ improvement. Both paths converge on the same liability-removing substitutions (D → A at position 27, N depletion at position 30) but diverge in their compensatory replacements, indicating that liability avoidance is orthogonal to the global hydrophobicity strategy, a separation enabled by motif-level reward design rather than whole-sequence property regression. The divergence arises because DPO pre-training encodes structural and expression-compatible preferences that constrain subsequent GDPO exploration to a hydrophobic-packing region of sequence space, whereas GDPO(SFT) must simultaneously learn liability avoidance and stability, resulting in a more conservative trajectory that retains polar residues. Despite the synthetic VHH library providing substantial (but non-uniform) combinatorial diversity at all degenerate CDR positions (8 in CDR1, 4 in CDR2, and 10 in CDR3), GDPO collapses this diversity at positions under strong biophysical constraint (positions 55–57 converge to 100% S-W-G consensus in both paths) while CDR3 positions 105–109 retain high entropy. Both trajectories therefore preserve a separation between developability-fixing and binding-flexible positions, and either path can be selected per epitope-specific requirements; both reduce liability and preserve humanness. A complementary mechanism is illustrated by IgGM, which achieves a similarly tight deamidation distribution (*σ* = 0.18, mean 5.01) by template-driven sampling rather than by reward optimization; the convergence of two distinct strategies on the same axis suggests that low-deamidation VHH design is a relatively constrained optimization problem rather than a unique GDPO contribution, while the wider competitor pool (*σ* = 1.6–3.2) shows that the constraint is not trivially captured by sequence priors alone.

### Practical implications

The reward-decomposition strategy used here (targeting specific sequence motifs and structural features grounded in established biophysical mechanisms rather than training end-to-end property predictors) is portable to other protein engineering tasks. Enzyme design could target catalytic-residue geometry and core-packing rewards as proxies for *k*_cat_ and stability; peptide therapeutics could optimize interface contacts and helicity as proxies for binding and pharmacokinetics; and de novo protein design could balance secondary-structure formation and binding-motif placement through analogous mechanistic decompositions. We propose, on the basis of these results, that multi-objective protein optimization may not require exhaustive property prediction for every desired attribute: a causal decomposition of fitness into independently optimizable, biophysically interpretable reward signals, together with an alignment algorithm that preserves each reward’s individual gradient, appears sufficient to capture the emergent biophysical gains we observe here. Whether this principle generalizes to other protein classes is an empirical question to be tested by future work.

### Limitations

Several limitations bear on the interpretation of these results.

#### In silico evaluation only and predictor overlap

All developability outcomes reported here are predictions from published sequence-based predictors (TEMPRO for T_*m*_ [45], NetSolP for solubility [42], Sapiens for humanness [41], instability index from amino-acid composition [43], and motif-based liability scores). Three of these predictors (TEMPRO, NetSolP, Sapiens) also enter the DPO composite preference score, so improvements on those axes are partly self-consistent; the GDPO rewards are sequence-motif and physicochemical features distinct from these predictors, but share inputs (amino-acid composition, hydrophobicity) with several. No experimental measurements of thermal stability, expression yield, aggregation, or binding are reported for the generated sequences; independent scorers (nanobody-specific nativeness tools, nanobody-calibrated developability profilers) and in vitro validation on a subset of generated candidates are essential before any therapeutic claim can be made and are outside the scope of this study.

#### Target scope

The 65 targets used here span linear peptide epitopes, intrinsically disordered protein fragments and whole folded protein domains, but the conditioning input to the model is always the linear sequence of the target; structural context is not provided to the generator. Generalization to highly conformation-dependent targets or to epitopes with non-canonical amino acids is therefore not tested by this study.

#### Novelty extent

Generated sequences differed from the nearest training sequence by a mean of 8.1–9.0 amino acids out of 126 at *T* = 0.7 (see Supplementary Information). This represents local variation within the training manifold rather than unconstrained de novo design. Novelty at higher sampling temperatures is reported in the Supplementary Information.

#### Dataset provenance

Training data were generated on a proprietary mRNA-display platform [46] using a synthetic VHH library [47]. The signal-to-noise ratio of the screening assay is not fully characterized in this work and may influence the quality of the supervision signal.

#### Humanness is inherited from the training scaffold

The landscape benchmark (Fig. 5) shows that NanoAbLLaMA, IgLM, and nanoBERT achieve higher Sapiens humanness than Aiki-GeNano by ∼0.05–0.09 (several population *σ*). This is a direct consequence of the training library used by Aiki-GeNano: every SFT, DPO-preferred and DPO-dispreferred sequence derives from the Contreras *et al*. [47] synthetic VHH library, which is built on the cAbBCII10 camelid VHH backbone of Conrath *et al*. [44] (NBv1) rather than a humanized antibody backbone. The DPO preference pairs therefore vary only within the randomized CDR positions of NBv1 (8, 4, and 10 degenerate residues in CDR1, CDR2, and CDR3); the framework was effectively constant across every comparison the model saw. Under this training distribution the DPO objective could only learn to prefer CDR-level chemical and thermal-stability features, not framework humanization, because no examples of better-or-worse frameworks were ever paired. GDPO similarly rewards sequence-intrinsic liabilities and VHH scaffold hallmarks that are structurally consistent with camelid framework residues. The Sapiens humanness scores therefore reflect the camelid-VHH nature of the NBv1 scaffold rather than any behavior acquired during Aiki-GeNano training. A natural next iteration is to widen the training pool to include humanized VHH scaffolds, or to add an explicit humanness reward that operates at framework positions; both extensions would require preference data outside the NBv1 library and are treated as future work.

#### Landscape benchmark scope

The head-to-head comparison in Fig. 5 was run on 10 GPCR target epitopes at seed 42 only; Aiki-GeNano sequences are accepted at the fixed 126-residue NBv1 backbone the library is built on, and external tool sequences within the canonical 110–130 AA VHH length window (|Cys|≥2, canonical 20-AA alphabet). Three further confounders bear on how the per-tool numbers should be interpreted. *(i) Scaffold lock-in*. The Aiki training data fixes 103 of the 126 sequence positions; only the 8 / 4 / 10 degenerate CDR1 / CDR2 / CDR3 positions of the NBv1 scaffold vary at generation time, whereas competitors emit across more diverse frameworks (NanoAbLLaMA varies ∼100 of 126 positions per output). The Aiki-GeNano family’s per-axis means therefore reflect optimization *within* one handpicked scaffold; competitor means are framework-averaged. *(ii) Per-tool sample size*.

Aiki contributes *N* ≈ 6,400 gated sequences per stage versus ∼ 1,000 per competitor. Standard errors on the mean are small on both sides (*<* 0.2 ^◦^C for *T*_*m*_), so the headline rankings are not changed by sample-size disparity; however, the per-tool *σ* widths in Fig. 5a should be read as framework- and CDR-diversity differences across tools rather than as predictor noise. *(iii) Canonical-VHH content of competitor outputs*. A stricter VHH-shape filter (canonical FR4 motif and N-terminal Q) classifies 100% of nanoBERT and IgGM outputs, 85% of NanoAbLLaMA, 91% of IgLM (the failures concentrate at the N-terminal Q residue that IgLM elides under our prompting), and only 0.6% of ProteinDPO as canonical VHHs. ProteinDPO’s headline values therefore reflect structure-conditioned, inverse-folded sequences that pass the loose 110–130 AA gate rather than canonical-VHH output, which may also explain its wider predicted-*T*_*m*_ distribution; we therefore suggest that its instability-index and NetSolP-solubility leadership be read with the qualification that those values are computed on predominantly novel-protein outputs rather than on canonical nanobody outputs.

## 4 Materials and Methods

### 4.1 Training data and preference construction

We synthesized a nanobody library described in the literature [47] based on a fixed 126-residue camelid VHH backbone, with 8 degenerate positions in CDR1, 4 in CDR2, and 10 in CDR3, leading to a theoretical library diversity of 10^18^. We then screened this library against 65 target epitopes using a proprietary mRNA display-based platform [46]. The dataset comprises 1.35 million nanobody–epitope pairs across 65 unique target epitopes (Supplementary Fig. S1). This epitope diversity ensures model generalization across varied binding contexts, spanning short linear peptides, intrinsically disordered regions and full folded protein domains.

For SFT, the training set comprised 100,000 library-member sequences per target obtained from the raw mRNA-display screening data of [46].

For DPO, we constructed a preference dataset of 522,800 pairs by clustering the top 10,000 binders per target and pairing them according to binding-affinity similarity. Pairs with similar affinity (|ΔlogKd| *<* 0.75; 50%, *n* = 261,400) were ranked by a composite developability score:

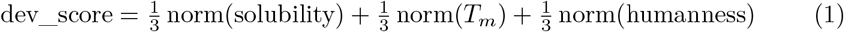

where solubility was predicted using NetSolP [42], thermal stability (*T*_*m*_) using TEMPRO [45], and humanness using the Sapiens antibody language model [41]. Pairs with divergent affinity (|ΔlogKd| ≥0.75; 50%, *n* = 261,400) were generated by treating binders of sequence-distant targets as non-binders. This partitioning prevents conflicting preference signals: when affinity differs substantially, binding dominates; when affinity is comparable, developability determines preference. The threshold of 0.75 log units was chosen to balance dataset partitioning (equal 50/50 split) while ensuring sufficient affinity discrimination in the divergent subset.

For GDPO, sequences were generated online through rollout sampling conditioned on the same 65 target epitopes used during SFT training, with no pre-constructed preference pairs required. The three-stage optimization (SFT → DPO → GDPO) progressively refines the initial library distribution toward higher-quality regions of nanobody sequence space.

### 4.2 Model training

#### Supervised fine-tuning

We adapted ProtGPT2 (738M parameters) [18], pre-trained on UniRef50, for epitope-conditioned nanobody generation using QLoRA [48]. Training examples were formatted as ChatML [49] sequences, with the epitope provided as the prompt and the nanobody sequence as the response, both encoded in FASTA format.

The model was fine-tuned using a response-only cross-entropy objective:

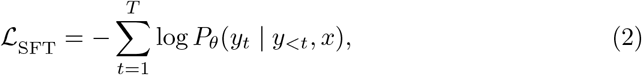

where *x* denotes the epitope and *y*_*t*_ the nanobody tokens. This stage establishes a binding-compatible sequence prior without explicit optimization for developability. Training was performed for 20,000 steps with an effective batch size of 48 on a single NVIDIA A100 40GB GPU (Supplementary Table S3).

#### Direct preference optimization

The SFT model was subsequently aligned using Direct Preference Optimization (DPO) [50] to bias generation toward more developable nanobody sequences. Training was performed on paired examples of preferred (*y*_*w*_) and dispreferred (*y*_*l*_) sequences using:

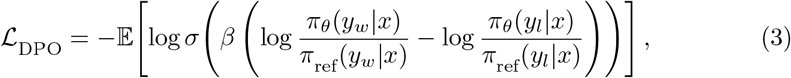

where *π*_ref_ denotes the frozen SFT model and *β* = 0.5 (Supplementary Table S4). Unlike prior applications of DPO in protein modeling, which are limited to small peptide datasets and single-property objectives [23], we scale to 1.35 million nanobody– epitope pairs across 65 targets and optimize a composite developability objective encompassing thermal stability, solubility, and humanness (Eq. 1).

#### Group reward-decoupled policy optimization (GDPO)

To enable simultaneous optimization of six independent mechanistic reward objectives without reward-signal collapse [35], we apply GDPO as a third training stage. For each epitope prompt, *G* = 16 nanobody candidates are sampled as rollouts, scored independently by each reward function, and used to update the policy. GDPO normalizes each reward *r*_*k*_ independently within its rollout group before aggregation:

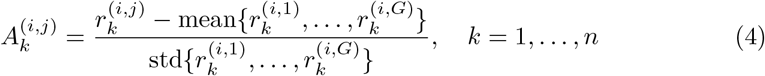

The weighted sum of per-reward advantages is then batch-normalized to stabilize the numerical scale as the number of rewards grows:

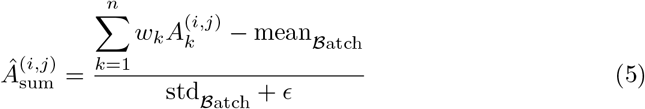

The policy is updated by maximizing a clipped surrogate objective with a KL divergence penalty, implemented as the BNPO (batch-normalised policy optimisation) loss in NVIDIA’s GDPO module of TRL:

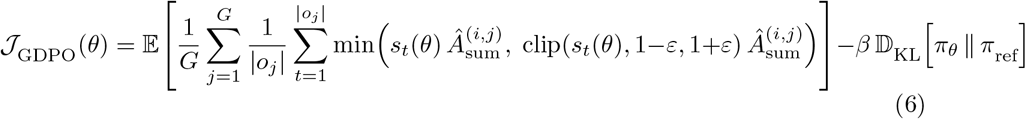

where 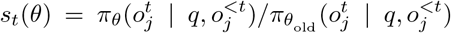 is the PPO-style importance ratio between the current and previous-iteration policy snapshots, ε = 0.1 is the clipping coefficient, and *β* = 10^−3^ is the KL penalty anchoring the policy to the frozen reference model *π*_ref_ (the SFT or DPO checkpoint, depending on the GDPO variant). The importance ratio and the KL anchor are therefore two distinct objects: the ratio compares against the rolling old-policy snapshot used for trust-region clipping, while the KL term compares against the frozen reference.

##### Algorithm 1

GDPO training (compact).

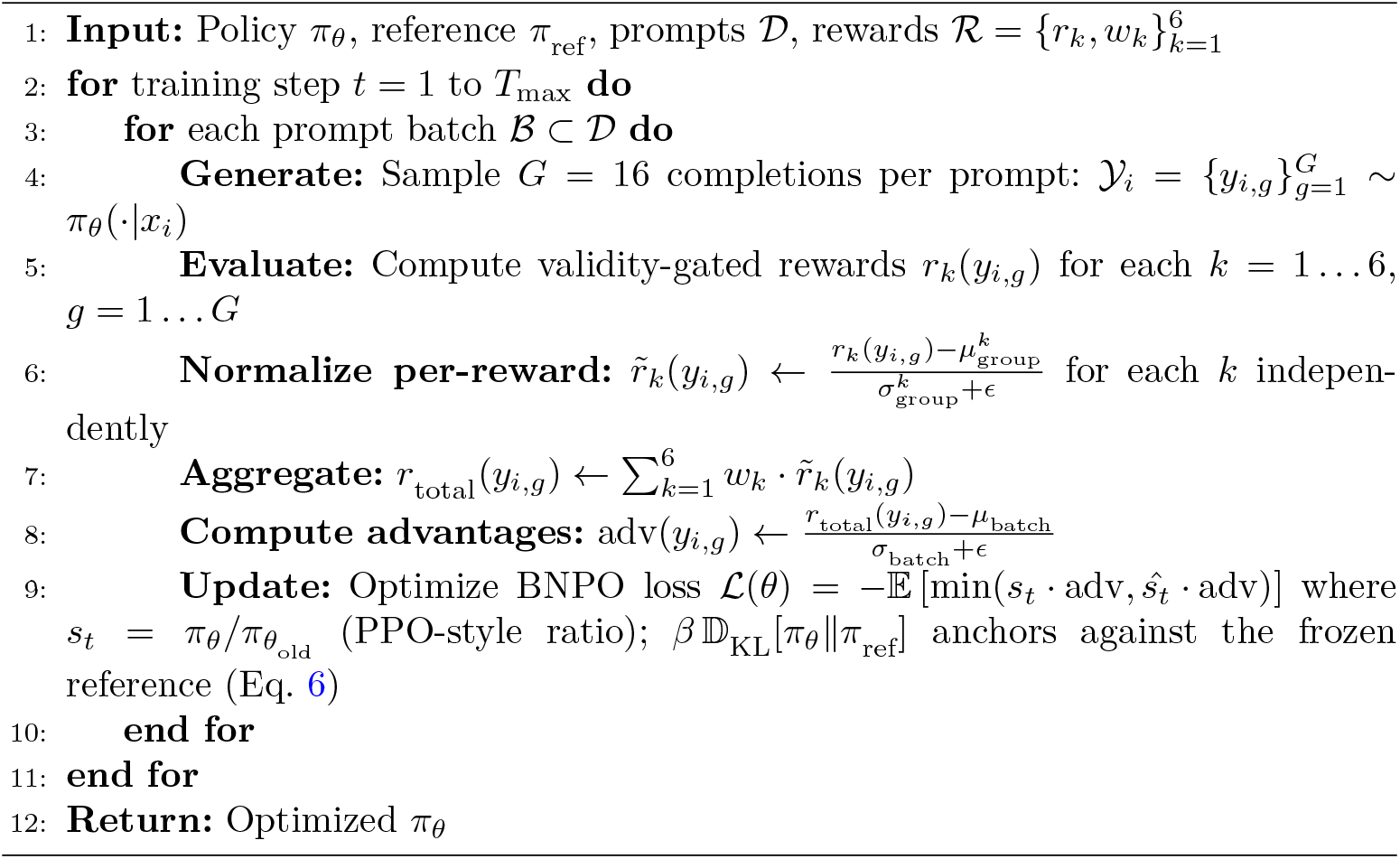

### 4.3 Reward design

We designed six sequence-based reward functions that collectively constrain the model to the VHH scaffold while driving it toward chemical developability. Three rewards directly penalize sequence features associated with poor drug developability, and three anchor the nanobody framework identity, ensuring the model cannot satisfy developability objectives by generating structurally invalid sequences (Table 1).

**Table 1.** GDPO reward functions and weights for therapeutic developability.

| Reward | Property | Weight | Rationale |
| --- | --- | --- | --- |
| $r_1$ | FR2 aggregation | 0.15 | VHH-specific solvent-exposed region |
| $r_2$ | Hydrophobic patches | 0.20 | Primary aggregation driver |
| $r_3$ | Chemical liability | 0.25 | Degradation motifs (highest priority) |
| $r_4$ | Expression probability | 0.15 | Manufacturability proxy |
| $r_5$ | VHH hallmark | 0.15 | Framework identity conservation |
| $r_6$ | Scaffold integrity | 0.10 | Structural validity (ungated) |

The FR2 aggregation reward (*r*_1_) computes mean residue hydrophobicity across FR2 (positions 36–53) using the Kyte–Doolittle scale [36], the established standard for identifying aggregation-prone regions in solvent-exposed domains. The hydrophobic-patch reward (*r*_2_) identifies consecutive runs of ≥ 5 residues from {I, L, V, F, M} (an approach inspired by AGGRESCAN [37, 51]) and penalizes their fractional coverage of the core sequence as a sequence-level aggregation-risk score. The chemical liability reward (*r*_3_) was designed from motif catalogues established in the antibody developability literature [52, 53]: deamidation (NG, NS), isomerization (DG, DS), fragmentation (DP), N-glycosylation (NxST), CDR methionine oxidation, and charge clusters are each assigned severity scores reflecting clinical impact, and total severity is mapped to [0, 1] via a hyperbolic function; this reward carries the highest weight, reflecting its direct relevance to regulatory and manufacturing requirements. The expression reward (*r*_4_) applies the Wilkinson–Harrison model [40], a predictor of soluble *E. coli* expression derived from turn-forming residue fraction and net charge balance. The VHH hallmark reward (*r*_5_) scores the four FR2 tetrad positions (Kabat 37, 44, 45, 47) based on residue classifications derived from Muyldermans [1]: VHH-canonical hydrophilic substitutions at these solvent-exposed positions are required due to the absence of a V_*L*_ pairing partner, and any reversion to hydrophobic VH-interface residues is penalized. The scaffold integrity reward (*r*_6_) is a continuous composite score enforcing exact 126-AA length, the C-terminal tag present in all library members, and the canonical Cys23–Cys104 disulfide bond [1]; this reward is deliberately ungated to maintain a persistent gradient signal throughout training.

A validity gate is applied to all five non-scaffold rewards. Sequences failing structural checks (incorrect length, missing C-terminal tag, non-standard amino acids, or fewer than two cysteines) receive zero reward across all gated channels:

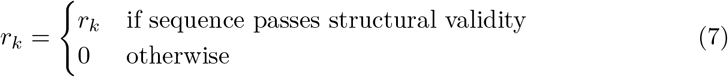

This gating is essential: without it, five reward channels provide meaningful signals for biologically invalid sequences, creating a gradient incentive to remain invalid. Ungated rewards also allow the model to exploit sequence length: generating truncated or oversized sequences that score favorably on individual reward channels without ever producing a valid nanobody, a form of reward hacking observed in early training runs. With the gate, the only path to non-zero reward is generating a structurally correct 126-AA VHH nanobody. The scaffold integrity reward (*r*_6_) is deliberately ungated, providing a persistent gradient signal that continuously pushes the model toward valid sequences even when all other reward channels return zero. Full reward calculations are provided in the Supplementary Information.

### 4.4 UMAP trajectory and landscape construction

For Supplementary Fig. S4, we drew a stratified subsample of 6,338 Aiki-GeNano-family sequences (25 per training-set target ×4 training-stage models, seed 42) and embedded each sequence by mean-pooling the last hidden state of *frozen* ProtGPT2 base `nferruz/ProtGPT2`, 738 M parameters; SFT-merged weights are deliberately not used, so the geometry is fair to all stages). UMAP was fit on the resulting embeddings (cosine metric, *n*_neighbors_ = 15, *d*_min_ = 0.05, 500 epochs, seed 42). Per-target trajectory threads in Fig. S4a include only targets where every stage has ≥5 post-gate sequences; 62 of 65 training-set targets meet this gate.

### 4.5 Reproducibility and compute

All training runs were performed on a single NVIDIA A100 40 GB GPU with a fixed random seed of 42 (Supplementary Tables S3–S5). SFT was performed for 20,000 steps, DPO for 6,000 steps, and each GDPO variant for 2,000 steps with *G* = 16 rollouts per prompt. For evaluation, three sampling seeds (42, 123, 456) and three sampling temperatures (*T* = 0.5, 0.7, 0.9) were used to compute the *n* = 3-seed robustness statistics in Results and in Supplementary Fig. S2; the head-to-head landscape benchmark in Fig. 5 uses seed 42 only. Training data were tokenized with the Prot-GPT2 tokenizer; sequences were formatted as ChatML messages [49] with the target epitope as prompt and the VHH as response. All property predictors (TEMPRO [45], NetSolP [42], Sapiens [41]) were run locally at published default settings. Sampling for evaluation used temperature *T* = 0.7 and nucleus *p* = 0.9 with 100 sequences per target; robustness to seed and temperature is reported in Supplementary Fig. S2. Reward-function source code is released with the training code (see *Code availability*). The training environment uses NVIDIA’s GDPO extension to TRL 0.18.0 (commit pinned in the released container image).

### 4.6 Landscape benchmark against contemporary generators

For the landscape comparison in Fig. 5, seven publicly available generators were installed and run on a shared panel of 10 GPCR target epitopes (UniProt accessions P41597, P21554, P32302, Q9H3N8, P35372, P61073, P49682, P51677, P41145, P25024). The five VHH-class head-to-head tools were nanoBERT [29] (masked-LM infilling of CDR positions on an NBv1 template); IgLM [54] (camelid-conditioned sampling); NanoAbLLaMA [30] (LLaMA2-7B with a single canonical camelid VHH N-terminal prefix `Seq=<QVQLV` with no explicit germline tag); ProteinDPO [24] (structure-conditioned inverse folding on a canonical VHH backbone); and IgGM [31] (diffusion + consistency model on an epitope-predicted complex). NanoAbLLaMA’s 6% length-tail beyond 130 AA is a configuration artefact of the generation wrapper rather than a property of the model; an explicit stop-token boundary on the `Seq=<…>` block would tighten validity to ∼99% without changing the per-sequence developability metrics for the 94% of outputs that already fall in-range. Two further tools, ProtGPT2 [18] and PepMLM [20], were included in the generation sweep for completeness but are not VHH-class generators under the prompting choices used here and are excluded from the head-to-head comparison. ProtGPT2 was prompted with an epitope prefix at the default sampling settings used for the other tools; in that configuration it returns median 20-AA outputs and 2/900 fall within the 110– 130 AA nanobody window, consistent with the model being a general-protein LM trained on UniRef50 rather than a VHH generator. ProtGPT2 has been shown to produce VHH-length sequences when seeded with the canonical QVQLVES opening, but that prompting strategy carries no epitope conditioning and would duplicate IgLM’s role as an unconditioned VHH baseline already present in the panel and was therefore not pursued. PepMLM was run at its native 10-AA peptide-binder output length; the model is trained on peptides only, and lengthening the mask token count is not a route to VHH-shaped output. Each tool generated 100 sequences per target at sampling temperature 0.7 (or the tool’s closest equivalent) on seed 42. Aiki-GeNano sequences were scored on the same developability pipeline as in §S.15. Aiki-GeNano sequences were accepted at the fixed 126-residue NBv1 backbone the library is built on (exact length, |Cys|≥ 2, canonical 20-AA alphabet); competitor sequences were accepted within the canonical 110–130 AA VHH length window (|Cys|≥ 2, canonical 20-AA alphabet). The filter asymmetry follows from Aiki using a fixed-length library scaffold while the other tools generate to broader length distributions. Effect sizes reported in Fig. 5a are computed as 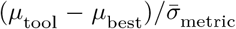, where 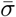 is the pooled cross-tool standard deviation for that metric. A 24-sequence predictor self-consistency check confirmed that the current NetSolP, TEMPRO, Sapiens, and motif-scoring implementations reproduce cached Aiki-GeNano values within Δ*T*_*m*_ ≤ 1 ^◦^C, ΔNetSolP ≤ 0.05, and exact motif match. The benchmark recipes (smoke-test scripts for each generator, patches applied to upstream codebases, and the per-tool aggregate metrics) are openly available at https://github.com/aikium-public/aiki-genano/tree/main/benchmarks, and the per-tool aggregate metrics file underlying Fig. 5 is reproduced in the Zenodo deposit (https://doi.org/10.5281/zenodo.19757842, file figure_data/fig5_landscape_benchmark.csv).

## Data availability

The nanobody–epitope screening dataset (1,354,410 pairs across 65 targets) generated on Aikium’s mRNA-display platform is proprietary and contains commercially sensitive sequences. The trained model checkpoints (SFT, DPO, GDPO (from DPO), GDPO (from SFT)) and the generated nanobody amino-acid sequences reported in this study are likewise proprietary.

The numerical data underlying the figures and tables in this study (computed developability properties — TEMPRO *T*_*m*_, NetSolP solubility, Sapiens humanness, biophysical descriptors, motif counts, and the six GDPO reward scores), organised both as per-figure aggregates over all 65 evaluated targets and as per-sequence property tables for the 10 representative GPCR targets disclosed in this study, plus the per-tool head-to-head benchmark table underlying Fig. 5, are deposited at Zenodo (https://doi.org/10.5281/zenodo.19757842) under CC-BY-NC-4.0.

## Code availability

The complete source code for the SFT, DPO, and GDPO training stages, the six reward-function implementations (*r*_1_–*r*_6_), the property-prediction evaluation pipeline (TEMPRO, NetSolP, Sapiens), the analysis notebooks that produce the paper figures, and the head-to-head benchmark recipes against the five contemporary VHH-class generators evaluated here (nanoBERT, IgLM, NanoAbLLaMA, ProteinDPO, IgGM) are openly available at https://github.com/aikium-public/aiki-genano under the MIT licence. A container image reproducing the training and inference environment (CUDA 12.1, PyTorch 2.2, transformers 4.57, NVIDIA’s TRL-GDPO fork at commit-pinned 0.18.0-gdpo with three compatibility patches applied at build time) is published at https://ghcr.io/aikium-public/aiki-genano:1.0.0; an interactive inference demo running the GDPO (from DPO) checkpoint is hosted at https://aikium--aiki-genano-fastapi-app.modal.run.

## Author contributions

V.M. conceived and supervised the study, designed the SFT and DPO training data, contributed to the landscape analysis, and edited the manuscript. J.D. implemented a preliminary SFT and DPO pipeline. R.S.M. adapted the pipeline to nanobodies, identified GDPO as the multi-reward alignment stage, designed the reward functions, performed all training, evaluation, and analyses, and wrote the manuscript. E.I. and S.S. designed and executed the experiments that generated the training set and contributed to manuscript review. All authors reviewed and approved the final manuscript.

## Acknowledgements

We thank the Synthetic Biology and Protein Sciences teams at Aikium Inc. for the mRNA-display screening campaigns that produced the nanobody–epitope binder pairs underlying this study, and the wider Aikium engineering team for the data-engineering infrastructure that made the SFT, DPO and GDPO training stages reproducible end-to-end. We thank the broader open-source community whose foundation models, predictors, and tools made this work possible: ProtGPT2 [18], the TRL preference-optimization library, AbNatiV [34], the Therapeutic Nanobody Profiler [33], TEMPRO [45], NetSolP [42] and the Sapiens antibody language model [41], the AGGRESCAN methodology and downstream hydrophobic-patch tooling [37, 51], and the contributors to the head-to-head generators we benchmark against (nanoBERT, IgLM, NanoAbLLaMA, ProteinDPO, IgGM). A portion of this work was enabled by cloud credits awarded by the Google for AI Startups program and by startup credits awarded by Modal for model hosting.

## Use of generative AI

Generative-AI assistants (Claude Opus from Anthropic, ChatGPT from OpenAI) were used to help draft, copy-edit and proof-read portions of this manuscript and to scaffold figure-generation and analysis scripts; all scientific claims, every numerical result, every figure caption and every algorithmic decision are the human authors’ responsibility. AI-generated suggestions were reviewed and edited by the authors prior to inclusion. Generative AI was not used for hypothesis generation, data synthesis, or interpretation of experimental or benchmark outcomes.

## Supplementary Information

### S.7 Dataset scale and epitope target diversity

**Fig. S1.**
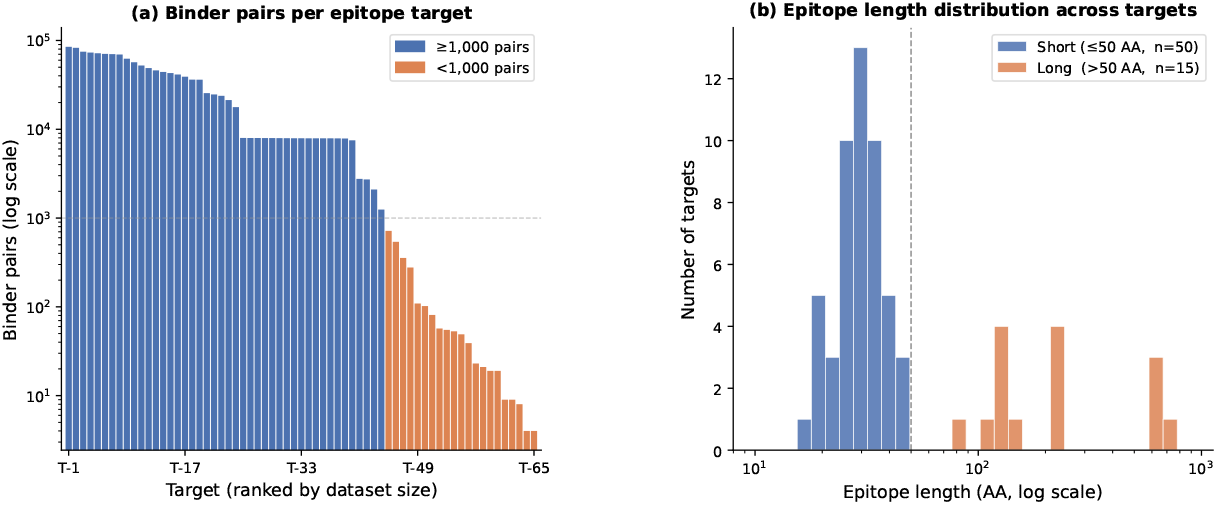
Dataset scale and epitope target diversity. **(a)** Distribution of nanobody–epitope binder pairs across 65 unique targets, ranked by dataset size (log scale; total 1,354,410 pairs). Targets with ≥1,000 pairs (*n* = 50) represent well-sampled antigen space after hierarchical clustering and deduplication; 15 targets contain *<*1,000 pairs. **(b)** Epitope length distribution across the 65 targets (log scale). The targets span three antigen classes: short linear peptides and intrinsically disordered region (IDR) fragments at the low-length end; whole folded protein domains in the mid range; and extended disordered regions or full IDPs at the high-length end. This breadth ensures that the language model is trained across both constrained linear epitopes and longer folded or flexible binding-relevant surfaces.

### S.8 Sequence novelty analysis

To confirm that generated sequences represent novel designs rather than memorization of training examples, we performed BLAST alignment of all generated sequences (seed 42, *T* = 0.7) against the full training dataset. Table S1 summarizes the similarity statistics for all four models. No model produces an exact match to any training sequence, indicating that generation is not memorising the training set. Mean percent identity to the closest training sequence ranges from 92.8% to 93.6%, corresponding to an average of 8.1–9.0 amino-acid differences across the 126-residue nanobody sequence. This indicates that while generated sequences preserve the overall nanobody fold and framework architecture, they incorporate substantial sequence variation relative to the training set. GDPO variants show slightly lower mean identity (92.8–93.1%) and higher mutation counts (8.7–9.0) than SFT and DPO baselines (93.4–93.6% identity, 8.1–8.3 mutations), consistent with reward-driven exploration beyond the training templates. GDPO(DPO), the model the paper calls Aiki-GeNano, sits at the bottom row of the table, matching its position in Fig. 5 and Table S2.

### S.9 Temperature-dependent sequence diversity patterns

Sampling temperature is a critical hyperparameter controlling stochasticity of sequence generation from language models. To understand how temperature affects the spatial distribution of sequence variability, we analyzed Shannon entropy profiles at three temperatures: *T* = 0.7, *T* = 0.9, and *T* = 1.2 (Supplementary Fig. S2c–d). At *T* = 0.7 (Fig. 3a, main text), all models exhibit high entropy exclusively in CDR regions, while framework regions remain nearly invariant. At *T* = 0.9, framework entropy increases modestly, particularly at positions 10–23, with DPO showing the strongest effect. At *T* = 1.2, framework variability increases further, with multiple positions outside CDRs exhibiting *H >* 1.0; GDPO(SFT) reaches *H >* 3.5 at CDR3 positions. For applications requiring strict nanobody scaffold preservation, *T* = 0.7 is therefore recommended; higher temperatures enable broader framework exploration at the cost of reduced structural predictability.

**Table S1.** Sequence similarity to training data via BLAST alignment.

| Model | Exact match (%) | Mean % identity | Mean mutations |
| --- | --- | --- | --- |
| SFT | 0.0 | 93.4 | 8.3 |
| DPO | 0.0 | 93.6 | 8.1 |
| GDPO(SFT) | 0.0 | 92.8 | 9.0 |
| GDPO(DPO) | 0.0 | 93.1 | 8.7 |

### S.10 CDR positional frequency analysis

Detailed positional analysis across 37 CDR positions reveals the spatial distribution of GDPO-driven modifications. Framework-proximal positions exhibit strict conservation across all models: CDR1 positions 24–26 (A-S-G), 32 (Y), 35 (F); CDR2 positions 48–54 (R-E-A-V-A-A-I), 58–59, 61–64 (T-Y-Y-A); and CDR3 positions 96– 101 (I-Y-Y-C-A-A), 112–114 (W-G-Q) all show 100% consensus. The strongest GDPO convergence occurs at three position classes: (i) liability hotspots (CDR1 position 27: D→A, 24%→2%, Δ*H* = −2.1 bits; position 30: N→I/F or Y, path-dependent, Δ*H* = −0.7 bits); (ii) structural hinges (CDR2 positions 55–57: S-W-G motif, 41– 63%→100%, Δ*H* = −1.8 bits each); and (iii) hypervariable loop anchors (CDR3 position 102: L conservation, 95%→98% in GDPO(SFT) vs. 77% in GDPO(DPO)).

### S.11 Three-way developability summary

We observed three concurrent shifts among Aiki-GeNano (GDPO(DPO)) sequences relative to the SFT baseline: chemical liability severity dropped (deamidation 6.8 → 5.4, isomerization 4.7 → 2.0; main-text Fig. 2f), predicted NetSolP solubility rose modestly (+0.024 on a 0–1 scale), and mean sequence hydrophobicity shifted toward less-hydrophilic GRAVY (−0.36 → −0.29) as liability-prone hydrophilic motifs (NG, NS, DG, DS) were removed. The three shifts co-occurred without any pair-wise penalty: the hydrophobicity move did not depress predicted solubility under NetSolP, and the liability drop did not depress humanness (Fig. 2c). A descriptive ellipse-on-ellipse view of these three axes was rendered during analysis but is not reproduced here because the four-model overlap of 1.5*σ* ellipses obscured rather than revealed the trade-off; per-tool numbers are tabulated in Supplementary Table S2.

**Fig. S2.**
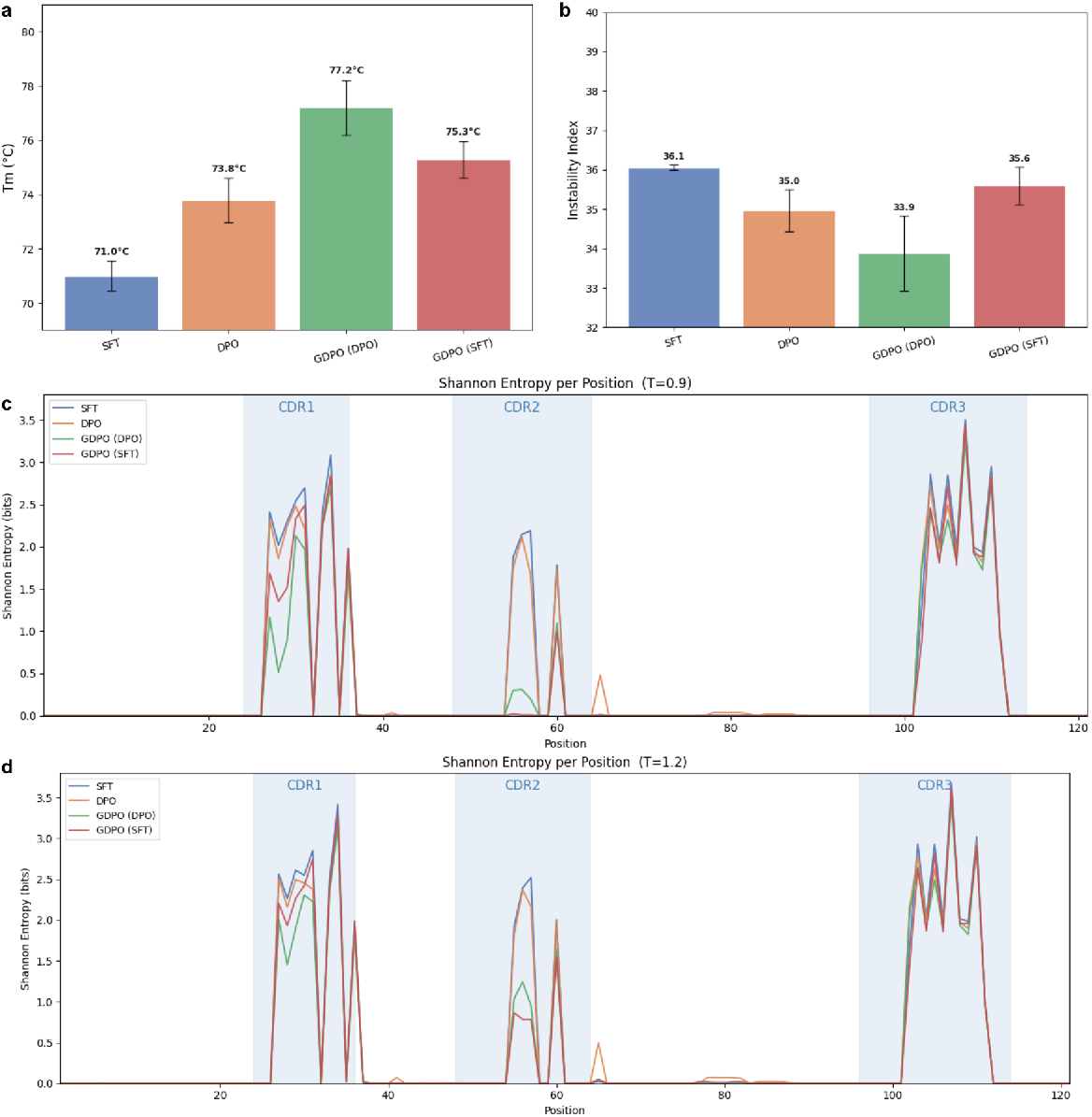
Property robustness and temperature-dependent sequence diversity. **(a**,**b)** Mean predicted T_*m*_ and instability index across all 9 generation runs (3 seeds ×*◦* 3 temperatures: *T* = 0.5, 0.7, 0.9). GDPO(DPO) consistently achieves the highest *T*_*m*_ (77.2 ± 1.0 °C, +6.2 ^*◦*^C over SFT) and lowest instability (33.9 ± 0.2) across all conditions. Model ranking remains stable regardless of seed or temperature. **(c**,**d)** Shannon entropy profiles at higher sampling temperatures *T* = 0.9 and *T* = 1.2 (seed 42).

### S.12 Landscape benchmark: supporting table and figures

The two Pareto trade-offs implied by Table S2 are: (i) Sapiens humanness vs. predicted *T*_*m*_, where NanoAbLLaMA wins humanness (0.750) and GDPO(DPO) wins *T*_*m*_ (78.3 ^◦^C) with no tool occupying both corners; and (ii) isomerization severity vs. *T*_*m*_, where the GDPO variants occupy the low-liability / high-*T*_*m*_ corner alone (Aiki isomerization *σ* ≈ 0.2–0.4 vs. 1.2–3.2 across the competitor pool). ProteinDPO’s predicted-*T*_*m*_ distribution is anomalously wide (*σ* = 9.4 ^◦^C vs. 3.6–5.9 for the other tools), consistent with its structure-conditioned outputs falling outside TEMPRO’s training distribution; the raw mean for that tool should be read with that caveat.

Output lengths from the five VHH-class tools concentrate within the 110–130 AA nanobody window (medians 121–127 AA), so the inclusion filter applied to competitor sequences does not exclude any substantial fraction of in-class output. ProtGPT2 (general-protein LM) returns median 20 AA outputs under our epitope-prefix prompting and PepMLM is a 10 AA peptide-binder generator; both are excluded from the head-to-head benchmark on length grounds and reported only in context (Methods §*Landscape benchmark against contemporary generators*).

**Fig. S3.**
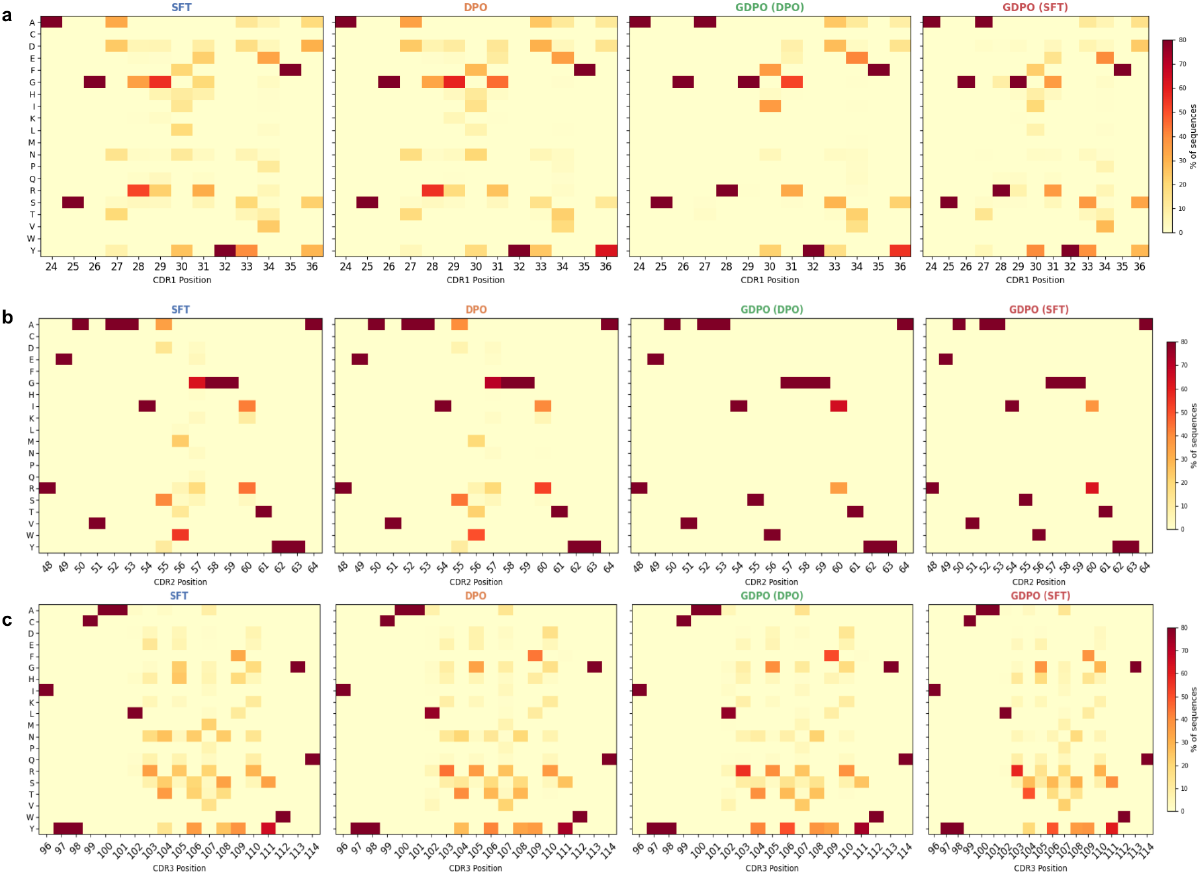
Amino-acid frequency heatmaps across CDR regions. **a–c**, Per-position amino-acid frequencies for CDR1 (a), CDR2 (b), and CDR3 (c) at *T* = 0.7 (all seeds). Each panel shows four model heatmaps (SFT, DPO, GDPO(DPO), GDPO(SFT)). Rows: 20 amino acids (grouped by properties); columns: CDR positions. Framework-proximal positions (CDR1: 24–26, 32, 35; CDR2: 48–54, 58–64; CDR3: 96–100, 112–114) show strong conservation (dark red bands) across all models. High-variability positions (CDR1: 27–31; CDR2: 55–57, 60; CDR3: 102–111) exhibit diverse distributions in SFT/DPO and targeted convergence in GDPO variants.

**Table S2.** Landscape benchmark headline table. Per-tool mean ±*σ* across all generated sequences accepted into the comparison, at seed 42 and sampling temperature *T* = 0.7. Aiki-GeNano sequences are accepted at the fixed 126-residue VHH backbone the library is built on (NBv1; Conrath *et al*. 2001 [44]); the other tools’ sequences are accepted within the canonical 110–130 AA VHH length window. Best value per column shown in bold. PepMLM and ProtGPT2 sit outside the nanobody length range under their designed use cases and are omitted from this comparison; their length context is given as a text note in the preceding subsection. Among the four Aiki rows, GDPO(DPO) (last row, bolded) is the model the paper calls Aiki-GeNano; SFT, DPO and GDPO(SFT) are training-stage ablations that show how each stage contributes.

| Tool | $N_{\text{valid}}$ | $T_m$ ($^{\circ}\text{C}$) | Instab. | Humanness | NetSolP sol. | Deam. | Isom. |
| --- | --- | --- | --- | --- | --- | --- | --- |
| SFT | 6,398 | $71.6 \pm 4.2$ | $36.1 \pm 5.4$ | $0.648 \pm 0.007$ | $0.611 \pm 0.023$ | $6.86 \pm 2.04$ | $4.72 \pm 1.31$ |
| DPO | 5,998 | $74.8 \pm 4.1$ | $34.3 \pm 4.3$ | $0.655 \pm 0.008$ | $0.627 \pm 0.023$ | $6.69 \pm 2.00$ | $4.58 \pm 1.46$ |
| GDPO(SFT) | 6,399 | $75.9 \pm 3.8$ | $35.1 \pm 5.2$ | $0.653 \pm 0.006$ | $0.618 \pm 0.020$ | $5.29 \pm 0.81$ | **$2.02 \pm 0.24$** |
| nanoBERT | 1,000 | $65.9 \pm 5.7$ | $31.6 \pm 4.8$ | $0.706 \pm 0.008$ | $0.577 \pm 0.025$ | $6.36 \pm 1.71$ | $4.19 \pm 2.01$ |
| IgLM | 997 | $68.3 \pm 5.0$ | $34.1 \pm 5.4$ | $0.740 \pm 0.039$ | $0.587 \pm 0.030$ | $6.82 \pm 1.64$ | $7.18 \pm 2.64$ |
| NanoAbLLaMA | 933 | $73.2 \pm 5.8$ | $32.9 \pm 5.7$ | **$0.750 \pm 0.014$** | $0.598 \pm 0.028$ | $6.37 \pm 2.19$ | $4.56 \pm 1.16$ |
| ProteinDPO | 1,000 | $56.3 \pm 9.4$ | **$28.9 \pm 9.7$** | $0.591 \pm 0.015$ | 0.630 | $6.77 \pm 3.63$ | $6.17 \pm 3.22$ |
| IgGM | 1,000 | $71.1 \pm 4.8$ | $33.3 \pm 4.5$ | $0.705 \pm 0.008$ | 0.576 | **$5.01 \pm 0.18$** | $5.26 \pm 1.94$ |
| **GDPO(DPO)** | **6,386** | **$78.3 \pm 3.5$** | **$32.8 \pm 3.8$** | **$0.657 \pm 0.007$** | **$0.635 \pm 0.020$** | **$5.39 \pm 0.96$** | **$2.06 \pm 0.36$** |

### S.13 Trajectory and per-sequence reinforcement on developability axes

Supplementary Fig. S4 renders two complementary views of how the pipeline moves sequences through predicted-developability space. Panel **a** plots the per-target SFT → DPO → GDPO(DPO) trajectory in three-property space (isomerization severity, lower = safer; predicted *T*_*m*_, higher = better; NetSolP solubility, higher = better), one thread per target with the population-mean trajectory in bold black, extending main Fig. 4a’s two-axis scatter to a third axis. Panel **b** plots the same set of sequences at the per-sequence level on the Aiki-GeNano-family UMAP, with each dot coloured by its own predicted property; per-stage means stamped inside each cell climb (top row) or drop (bottom row) monotonically across the three stages, and the regions of embedding space the pipeline enriches are the same regions that hold high-*T*_*m*_ / low-isomerization sequences. Both colormaps are oriented so the developable end is the darker end (yellow = low *T*_*m*_ / low isomerization; dark green / dark red = high *T*_*m*_ / high isomerization). The GDPO(SFT) bypass route is omitted from both panels; it is covered in main-text §*Target-level generalization and path-dependent optimization*.

**Fig. S4.**
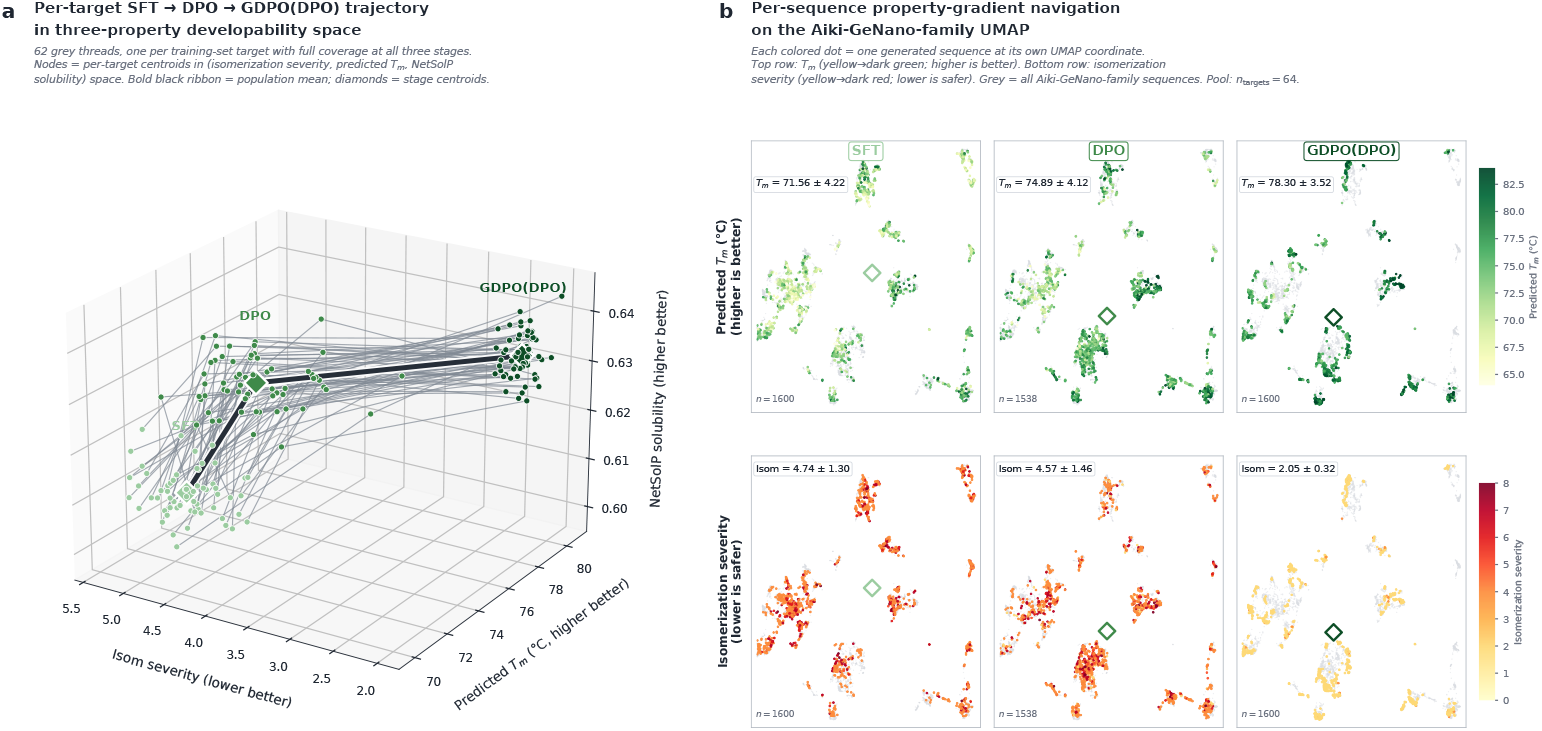
Trajectory in property space and per-sequence reinforcement on the Aiki-GeNano-family UMAP. **a**, Per-target SFT → DPO → GDPO(DPO) trajectory in three-property developability space. Each thin grey thread is one of 62 training-set targets with full coverage at all three stages (62 pass the ≥5 sequences/stage gate out of 65 trained targets); nodes are per-target centroids in (isomerization severity, predicted *T*_*m*_, NetSolP solubility) space; bold black ribbon = population-mean trajectory; diamonds = stage centroids. **b**, Per-sequence property-gradient navigation on the Aiki-GeNano-family UMAP. Top row: *T*_*m*_ (yellow = low, dark green = high; higher is better). Bottom row: isomerization severity (yellow = low, dark red = high; lower is safer). Each colored dot is one generated sequence at its own UMAP coordinate, colored by its own predicted property value; grey = all Aiki-GeNano-family sequences for context; diamonds = per-stage centroid. Per-cell *µ* ± *σ* and per-cell *n* stamped inside each cell; pool covers *n*_targets_ = 64 training-set targets with at least one Aiki-GeNano-family sequence in the gated subsample.

### S.14 Training hyperparameters

**Table S3.** SFT training hyperparameters.

| Hyperparameter | Value |
| --- | --- |
| Base model | ProtGPT2 |
| LoRA rank $r$ | 64 |
| LoRA alpha $\alpha$ | 128 |
| LoRA dropout | 0.05 |
| Quantization | 4-bit NF4 |
| Loss | Response-only cross-entropy |
| Max steps | 20,000 |
| Batch size (per device) | 12 |
| Gradient accum. steps | 4 |
| Effective batch size | 48 |
| Max sequence length | 1,024 |
| Optimizer | AdamW |
| Learning rate | $2 \times 10^{-4}$ |
| LR scheduler | Cosine |
| Warmup steps | 500 |
| Weight decay | 0.01 |
| Gradient clipping | 1.0 |

**Table S4.**
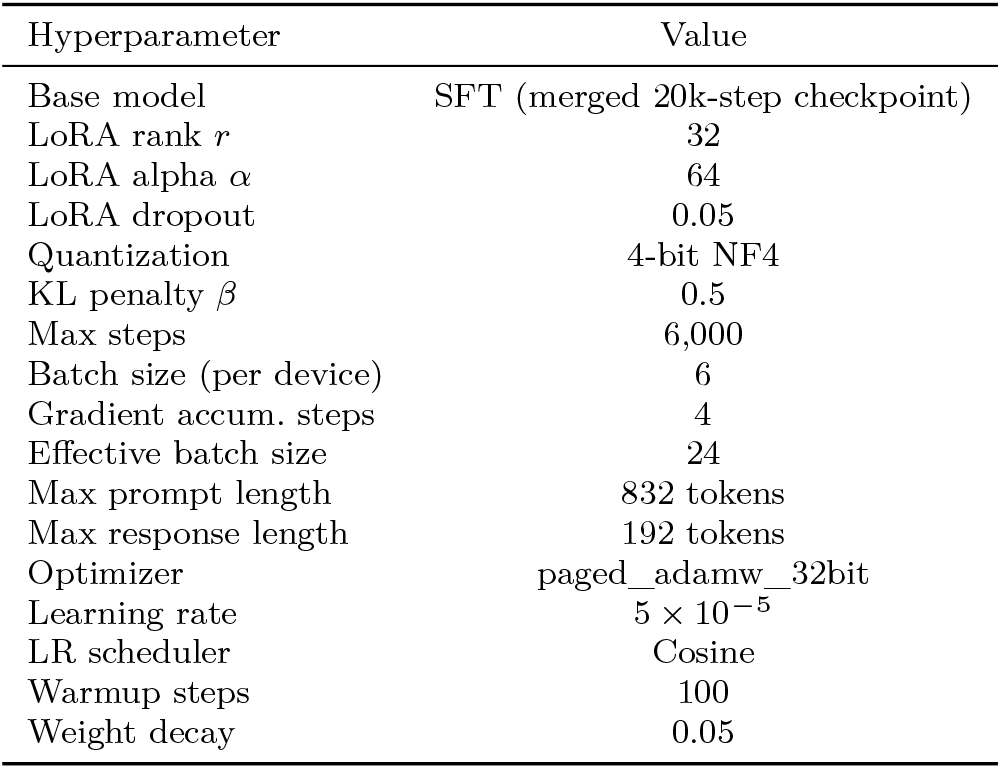
DPO training hyperparameters.

**Table S5.** GDPO training hyperparameters. GDPO(DPO) and GDPO(SFT) use identical settings; they differ only in initialization: GDPO(DPO) starts from the DPO checkpoint at step 6,000, GDPO(SFT) starts from the merged SFT model.

| Hyperparameter | Value |
| --- | --- |
| Initialization | DPO (step 6,000) *or* SFT (merged) |
| LoRA rank $r$ | 32 |
| LoRA alpha $\alpha$ | 64 |
| LoRA dropout | 0.05 |
| Precision | fp16 |
| Rollouts per prompt $G$ | 16 |
| Max completion length | 60 tokens |
| Sampling temperature | 0.9 |
| Top-$p$ (nucleus) | 0.92 |
| Top-$k$ | disabled |
| KL penalty $\beta$ | $1 \times 10^{-3}$ |
| Clip coefficient $\varepsilon$ | 0.1 |
| Loss type | BNPO |
| Disable dropout | Yes |
| Training steps | 2,000 |
| Batch size (per device) | 16 |
| Gradient accum. steps | 4 |
| Effective batch size | 64 |
| Optimizer | paged_adamw_32bit |
| Learning rate | $1 \times 10^{-5}$ |
| LR scheduler | Cosine |
| Warmup steps | 200 |
| Weight decay | 0.05 |
| Gradient clipping | 1.0 |
| Random seed | 42 |

### S.15 GDPO reward function definitions

All reward functions return values in [0, 1] where 1.0 indicates the optimal outcome. Rewards *r*_1_–*r*_5_ are validity-gated: any sequence failing the structural validity check receives 0.0 on all five channels. Reward *r*_6_ is ungated and evaluated on all sequences.

#### Structural validity gate

A sequence passes the validity check if and only if: (i) length = 126 AA; (ii) C-terminal tag; (iii) all residues belong to the standard 20-AA alphabet; and (iv) cysteine count ≥ 2.

#### r_5_: VHH hallmark (weight w = 0.15, gated)

Based on the FR2 tetrad classifications of Muyldermans [1]. For each of the four Kabat positions *p* ∈ {37, 44, 45, 47}, the residue *aa*_*p*_ in the core sequence (positions 1–121) is scored as:

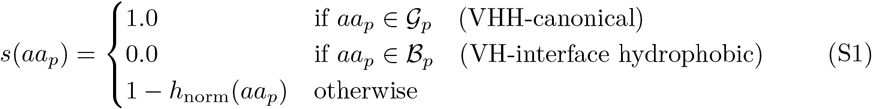

where 𝒢_*p*_ and ℬ_*p*_ are the canonical and penalized residue sets per position (Table S6), and *h*_norm_ is the min–max-normalized Kyte–Doolittle score [36]. The reward is the mean over all four positions:

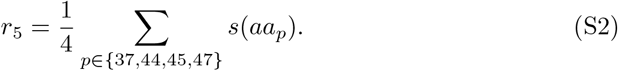

**Table S6.**
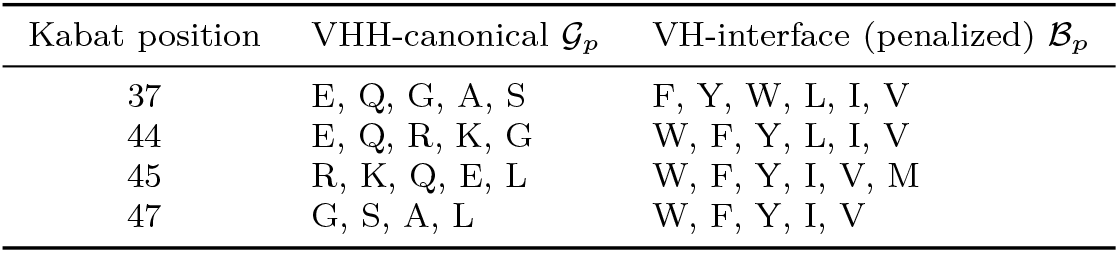
VHH hallmark residue classifications per FR2 tetrad position.

#### r_1_: FR2 aggregation (weight w = 0.15, gated)

Kyte–Doolittle hydrophobicity values [36] are min–max-normalized to [0, 1] over the full 20-AA alphabet (*h*_min_ = − 4.5 for Arg, *h*_max_ = 4.5 for Ile). Mean normalized hydrophobicity is computed over the FR2 region (core positions 36–53) and the reward penalizes high hydrophobicity:

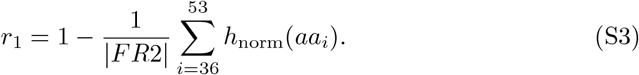

#### r_6_: scaffold integrity (weight w = 0.10, ungated)

A continuous composite score evaluated on the full 126-AA sequence (including the C-terminal tag):

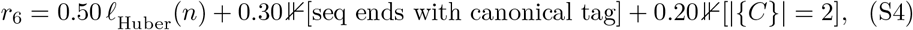

where *n* is sequence length and *ℓ*_Huber_ is a Huber-loss length-proximity score with *δ* = 2, target length *L*^∗^ = 126, and maximum deviation *d*_max_ = 10.

#### r_3_: chemical liability (weight w = 0.25, gated)

Motif scanning is performed on the core sequence (121 AA). Each motif occurrence contributes its severity score *s*_*m*_ to the total severity *S*. Motif definitions and severity scores are given in Table S7. Total severity is mapped to [0, 1] via a hyperbolic function with *k* = 10:

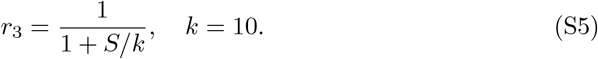

**Table S7.**
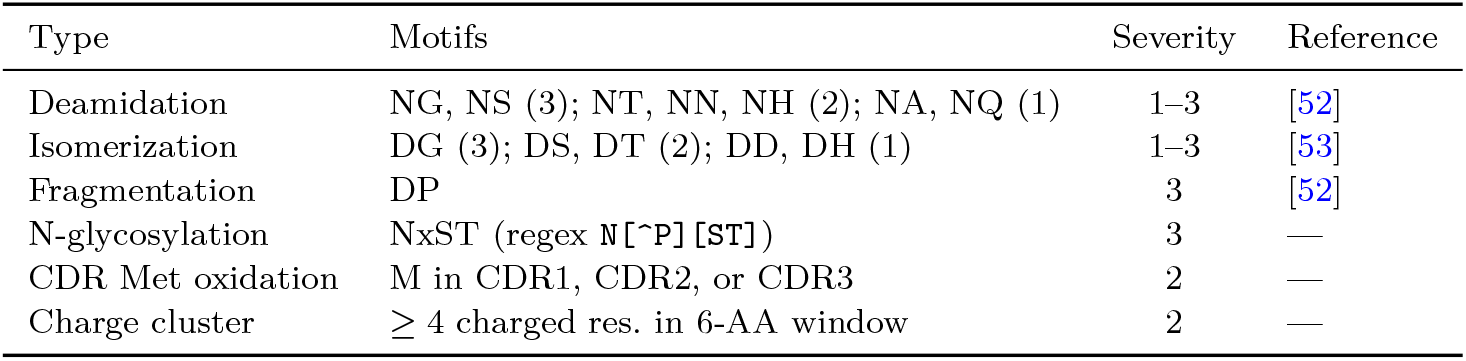
Chemical liability motifs and severity scores.

#### r_2_: Hydrophobic patches (weight w = 0.20, gated)

Consecutive runs of residues from ℋ = {I, L, V, F, M} of length ≥ *L*_min_ = 5 in the core sequence are identified as aggregation hotspots, inspired by the AGGRESCAN methodology [37, 51]. The reward penalizes fractional coverage:

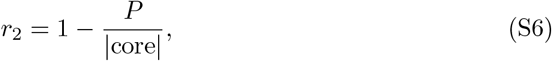

where *P* is the total number of residues in all patches and |core| = 121.

#### r_4_: expression probability (weight w = 0.15, gated)

The Wilkinson–Harrison model [40] predicts the probability of soluble *E. coli* expression from two sequence statistics computed over the core sequence: the turn-forming residue fraction *f*_turn_ over {N, G, P, S}, and the mean charge 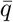 over {R, K} and {D, E}. The canonical CV score is:

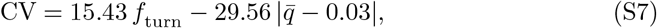

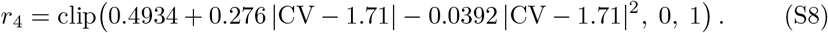

